# Conformational dynamics of the bacterial E3 ligase SspH1

**DOI:** 10.1101/2025.05.12.653404

**Authors:** Cassandra R. Kennedy, Diego Esposito, David House, Katrin Rittinger

## Abstract

The SspH/IpaH family of novel E3 ligases (NELs) are found in a number of Gram-negative bacteria and are used to target host enzymes for degradation to support pathogenesis. These E3 enzymes are autoinhibited in the absence of substrate and different models for release of autoinhibition have been suggested. However, many of the molecular details of individual steps during the ubiquitin transfer reaction remain unknown. Here, we present the crystal structure of *Salmonella* SspH1 and an analysis of the solution properties of SspH1 on its own and in complex with substrate and ubiquitin. Our data show that SspH1 exists in a conformational equilibrium between open and closed states and that substrate binding only modulates the distribution of these states but does not induce major conformational changes. This suggests that additional mechanisms must exist to bring the substrates close to the active site to mediate transfer of ubiquitin from the E3∼Ub conjugate.

## INTRODUCTION

Gram-negative bacteria remain a huge disease burden globally. *Shigella* and non-typhoidal *Salmonella* together account for an estimated 660,000 human deaths worldwide each year.^1^ A key part of *Salmonella* and *Shigella* pathogenesis is the delivery of virulence proteins (effectors) into the host cell through a Type 3 Secretion System (T3SS) to interfere with host immune responses and support bacterial replication.^2,3^ Effector proteins perform a range of cellular functions and often have enzymatic activities that include proteases, acetyltransferases, kinases, phosphatases, E3 ubiquitin ligases and deubiquitinases (DUBs).^2^ Ubiquitination plays an important role in the regulation of eukaryotic cellular processes and is a key mechanism to target proteins for proteasomal degradation. It is mediated by a 3-step enzymatic cascade including E1 activating, E2 conjugating and E3 ligase enzymes.^4^ Intriguingly, bacteria do not have a canonical ubiquitin system themselves but have evolved proteins that mimic and hijack the host ubiquitin system to support their survival and proliferation.^5–7^ Many pathogenic bacteria contain E3 ligases most of which structurally resemble their eukaryotic counterparts, yet some bacteria including *Salmonella* and *Shigella* have evolved a new class of E3 ligases, the Novel E3 Ligase family (NELs), that have no structural homology to eukaryotic E3s in their catalytic domain but function via an active site cysteine, analogous to HECT or RBR E3 ligases.^8,9^ These catalytic cysteine-containing E3 ligases transfer ubiquitin onto the substrate in a 2-step reaction: first ubiquitin is transferred from the E2∼Ub conjugate to form an E3∼Ub conjugate in a transthiolation reaction, and subsequently onto lysine residues in the substrate or ubiquitin itself via an aminolysis reaction.

NELs are found in a number of gram-negative bacteria such as the mammalian pathogens *Salmonella* and *Shigella*, and plant pathogens *Ensifer fredii* and *Ralstonia solanacearum*.^10^ They are composed of an N-terminal secretion motif, a leucine-rich repeat (LRR) domain and a C-terminal NEL catalytic domain and are structurally conserved across the family. Crystal structures and mechanistic studies of NELs have provided insight into the activities of the two key functional domains: the LRR domain,^9^ which is responsible for substrate recognition, and the NEL domain,^8^ which can be divided into two subdomains: the C-terminal subdomain (CSD) containing the E2-Ub binding ‘thumb’ (E2-UbBD) and the N-terminal subdomain (NSD) which encompasses the catalytic E3 region containing the active site cysteine.^11^ A linker region joins the LRR and NEL domains together, however the mechanistic role and importance of flexibility of this linker remains poorly defined.

The *Shigella* NEL family members comprise IpaH proteins, with the best characterized members being IpaH9.8, IpaH1.4/2.5 and IpaH7.8, whose targets include GBP1,^12–14^ Ste7,^15^ NEMO,^16^ LUBAC,^17^ and Gasdermin B.^18^ The *Salmonella* NEL subfamily includes SspH1, SspH2 and SlrP, which target PKN1,^19,20^ NOD1/SGT1,^21^ and thioredoxin,^22^ respectively. Upon ubiquitination, these host proteins are directed to proteasomal degradation, thereby suppressing the host immune and inflammatory response to bacterial infection.

The exact mechanism underlying the regulation of NEL E3 ligase activity has been the subject of many studies. The isolated NEL domain is constitutively active and forms free ubiquitin chains, while ubiquitination activity is suppressed in the full-length proteins.^8^ Auto-inhibition of NEL activity is important to prevent auto-ubiquitination and subsequent proteasomal degradation or formation of unanchored ubiquitin chains which could stimulate a host immune response.^23–26^ Initially, NEL proteins were thought to be auto-inhibited due to steric blockage of the catalytic region by the LRR domain, thereby preventing access to the E3 active site, until release of inhibition by substrate binding.^27^ This interpretation was based on crystal structures of NEL ligases including IpaH3,^9^ SspH2,^28^ and IpaH9.8,^29^ capturing two distinct ‘open’ and ‘closed’ conformations of the NEL proteins. In the ‘closed’ conformation (SspH2, PDB: 3G06), the LRR domain is oriented towards the NEL domain thereby apparently occluding access to the catalytic cysteine. In the ‘open’ conformations (IpaH3, PDB 3CVR; IpaH9,8, PDB 6LOL), the LRR domain is orientated away from the NEL domain, and these were assumed to represent active conformations. However, elegant biochemical experiments using mutants of the catalytic acid and base residues have since shown that NELs are not auto-inhibited but can form an E3∼Ub intermediate, which undergoes rapid non-productive ubiquitin turnover in the absence of substrate, indicating that the catalytic cysteine is accessible even in the absence of substrate binding.^30^ In addition, a recent study on IpaH9.8 has suggested that two residues in the concave surface of its LRR sense substrate binding to induce a conformational change and release a hydrophobic cluster that mediates auto-inhibition. This mechanism requires an amplifier loop in the LRR domain which however is not present in all family members.^29^

Currently it is unclear if these described regulatory mechanisms apply across the entire NEL family, nor have the intermediate steps in the ubiquitin transfer reaction been captured structurally. The first step in substrate ubiquitination, the initial transfer of ubiquitin from E2 to the NEL has been described to rely on significant intradomain flexibility between the catalytic E3 region and the E2-UbBD thumb^11^. In contrast, the structural determinants that mediate the switch from unproductive hydrolysis of the E3∼Ub intermediate to ubiquitin transfer onto substrate lysine residues is not understood. Similarly, the effect of substrate binding on the conformational flexibility of the LRR and NEL domains with respect to one another has not been studied.

To address these questions, we present a comparison of catalytic activities of NEL proteins across the *Shigella* and *Salmonella* subfamilies, highlighting similarities and differences between family members. To better understand the molecular mechanism of SspH1, we combine the structural characterization of SspH1 by crystallography with SAXS and NMR analysis to interrogate the solution state structure and conformational flexibility of SspH1 when bound to its substrate PKN1 and to ubiquitin. Our data indicate that the open and closed forms of SspH1 are in a conformational equilibrium and that substrate binding modulates the distribution between closed and open states but does not induce major conformational changes. Similarly, substrate binding has only a minor effect on the flexibility of ubiquitin in the E3∼Ub conjugate suggesting that additional as yet unknown mechanisms must exist to bring the LRR-bound substrate close to the ubiquitin thioester intermediate and initiate ubiquitin transfer.

## RESULTS

### Comparison of IpaH1.4, IpaH9.8, SspH1 and SspH2 catalytic activities

NELs are highly homologous in their catalytic NEL domain, and many residues previously identified as important for catalytic activity, including residues responsible for E2-Ub binding as well as catalytic acid and catalytic base residues, are conserved across the family. The main differences between NEL family members are in the LRR domains, which mediate substrate recognition but may also affect ubiquitin transfer as suggested for the amplifier loop in IpaH9.8. To investigate if the mechanism of ubiquitin transfer of NELs is conserved across the family we compared catalytic activities across 4 family members: two *Shigella*, IpaH1.4 and IpaH9.8, and two *Salmonella* proteins, SspH1 and SspH2, as these have well established substrates. A sequence alignment of these family members is shown in Supplementary Figure 1A,31 highlighting the conservation of key catalytic residues but also differences such as the lack of the IpaH9.8 amplifier loop in SspH1 and SspH2. The LRR domain of SspH2 is longer compared to the other three proteins, corresponding to 12 LRR repeats compared to 8 repeats for IpaH9.8 and 1.4.^28,29^ Structural models generated by AlphaFold2 predict an additional globular N-terminal domain in SspH1 and SspH2 which is structurally conserved across the two proteins (Supplementary Figure 1B),^32–34^ and found in other bacterial effectors such as SifA, though its function in E3 ligase activity has not been investigated thus far.

To interrogate the mechanisms of ubiquitin transfer across the *Shigella* IpaH and *Salmonella* SspH proteins, we purified three constructs for each of IpaH1.4, IpaH9.8, SspH1 and SspH2: the full-length protein (FL); the NEL domain (NEL); and the full-length protein minus the N-terminal secretion signal or globular domain in SspH1 and SspH2 (ΔN) (Figure 1A). First, we confirmed substrate ubiquitination activity of IpaH1.4, IpaH9.8 and SspH1 with their respective substrates LUBAC, GBP1 and PKN1 HR1b domain for the ΔN (Figure 1B) and FL constructs (Supplementary Figure 2A). Substrate ubiquitination by SspH2 could not be investigated due to difficulties with producing sufficiently pure SGT1/NOD1 substrate complex. For each construct of each protein, we performed a further three types of assays to assess activity: first we performed *in vitro* ubiquitination assays, where overall E3 ligase activity is monitored through autoubiquitination or formation of free ubiquitin chains. Next, we carried out E2∼Ub complex discharge assays using fluorescently labelled ubiquitin to isolate the E2∼Ub and E3∼Ub formation steps of the ubiquitin cascade. If a stable E3∼Ub intermediate is formed this can be observed on non-reducing SDS gels. Finally, we performed ubiquitin-loading assays with a covalent ubiquitin probe (ubiquitin vinyl sulfone, UbVS) to assess accessibility of the E3 catalytic cysteine.

**Figure 1.**
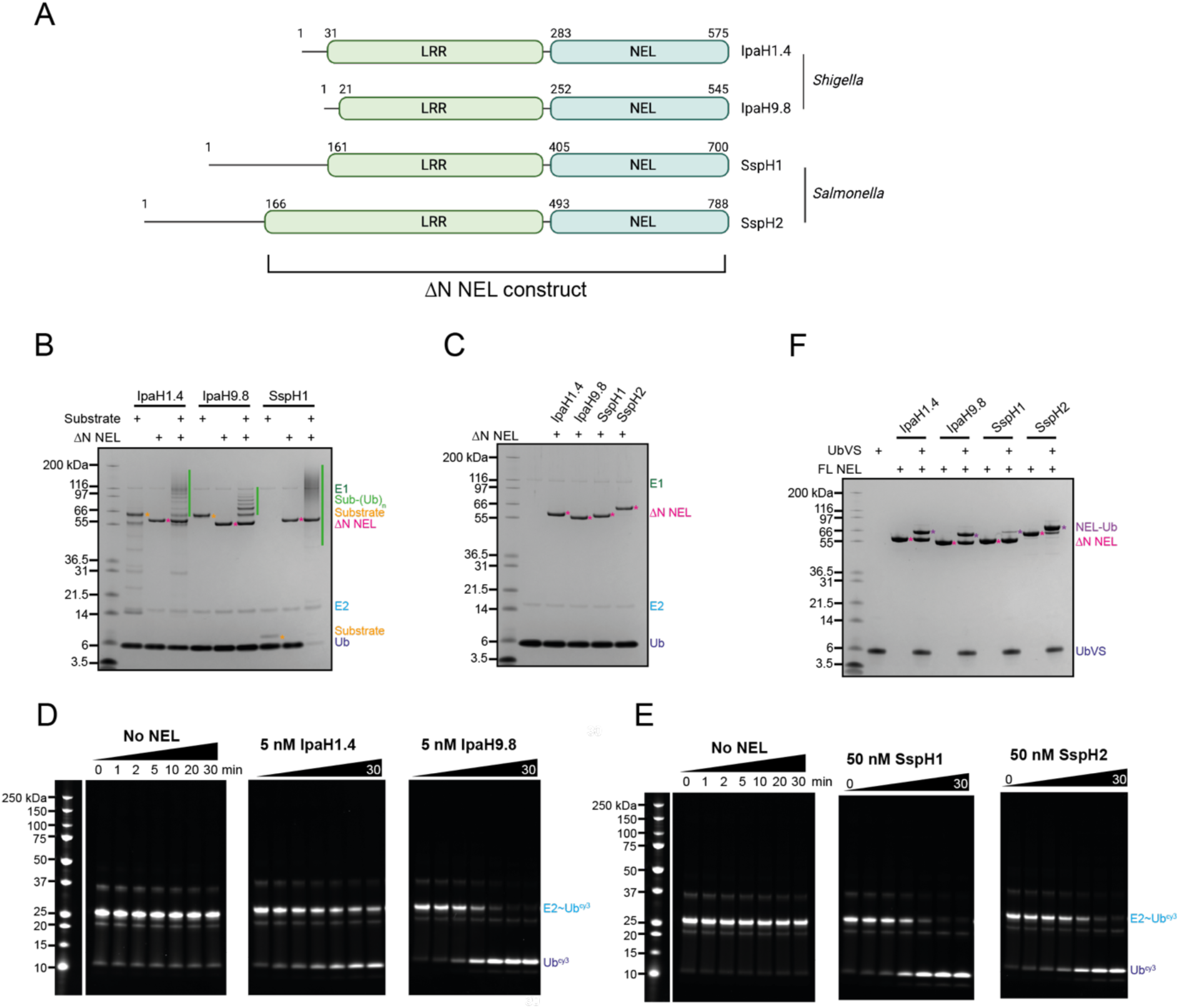
ΔN NEL activity assays. A) Diagram of constructs of NEL bacterial E3 ligases IpaH1.4, IpaH9.8, SspH1 and SspH2; B) Substrate ubiquitination assay with ΔN constructs of IpaH1.4 (substrate LUBAC), IpaH9.8 (substrate GBP1) and SspH1 (substrate HR1b PKN1). Performed with 0.1 µM UBA1 (E1), 2 µM UbcH5A (E2), 1.0 µM ΔN NEL (E3), 20 µM ubiquitin, 10 mM ATP at RT for 30 minutes with either 1 µM (LUBAC, GBP1) or 2 µM (HR1b PKN1) substrate. C) Auto-ubiquitination assay with ΔN constructs of IpaH1.4, IpaH9.8, SspH1 and SspH2. Performed with 0.1 µM UBA1 (E1), 2 µM UbcH5A (E2), 1 µM ΔN NEL (E3), 20 µM ubiquitin, 10 mM ATP at RT for 30 minutes. D) E2∼Ub discharge assay with ΔN IpaH1.4 and IpaH9.8. Performed with 1 µM UbcH5A∼Ub-cy3 (E2∼Ub^cy3^), 5 nM NEL (E3) at RT for 0-30 minutes. E) E2∼Ub discharge assay with ΔN SspH1 and SspH2. Performed with 1 µM UbcH5A∼Ub-cy3 (E2∼Ub^cy3^), 50 nM NEL (E3) at RT for 0-30 minutes. Assays for full length constructs are in Supplementary Figure 2. F) Ubiquitin-loading assay with ΔN constructs of IpaH1.4, IpaH9.8, SspH1 and SspH2. Performed with 20 µM UbVS, 5 µM ΔN NEL (E3) at RT for 2 hours.

As expected, we observe some similarities in activities across the four *Shigella* and *Salmonella* NELs. In substrate ubiquitination assays, we observed equal substrate ubiquitination with both ΔN and FL proteins indicating that the N-terminal globular domain present in SspH1/2 has no effect on catalytic activity (Figure 1B, Supplementary Figure 2A, Supplementary Figure 3A-C). In contrast, in the absence of substrate, no auto-ubiquitination or free chain formation activity was observed with ΔN and FL proteins (Figure 1C, Supplementary Figure 2B). In E2∼Ub discharge assays, FL and ΔN constructs discharge ubiquitin from UbcH5A∼Ub (Figure 1D and E, Supplementary Figures 2C and 3E) indicating that their catalytic cysteine is accessible but no stable E3∼Ub thioester intermediate is formed. This is in agreement with the labelling observed with UbVS in ubiquitin-loading assays (Figure 1F, Supplementary Figure 2D). The isolated NEL domains alone did not auto-ubiquitinate, however, free ubiquitin chains were synthesized indicating that removal of the LRR domain released auto-inhibition (Figure 2A, Supplementary Figure 3D). Similarly, in E2∼Ub discharge assays with isolated NEL domains some free di-ubiquitin was synthesized (Figure 2B), and all constructs were labelled by UbVS (Figure 2C).

**Figure 2.**
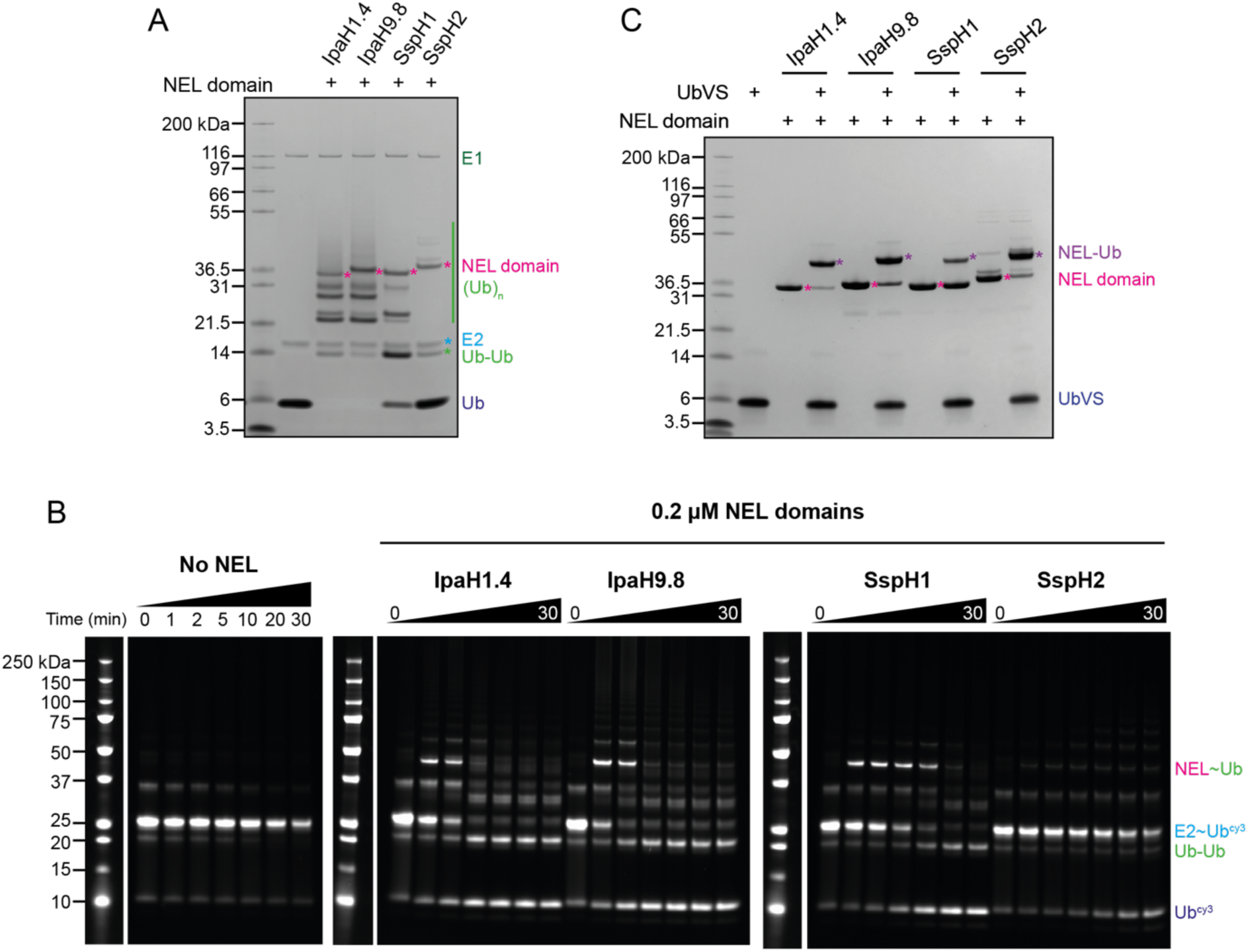
NEL domain activity assays. A) Auto-ubiquitination assay with NEL domains of IpaH1.4, IpaH9.8, SspH1 and SspH2. Performed with 0.1 µM UBA1 (E1), 2 µM UbcH5A (E2), 1 µM NEL domains (E3), 20 µM ubiquitin, 10 mM ATP at RT for 30 minutes. B) E2∼Ub discharge assay with FL constructs of IpaH1.4, IpaH9.8, SspH1 and SspH2. Performed with 1 µM UbcH5A∼Ub-cy3 (E2∼Ub^cy3^), 0.2 µM NEL domain (E3) at RT for 0-30 minutes. C) Ubiquitin-loading assay with NEL domains of IpaH1.4, IpaH9.8, SspH1 and SspH2. Performed with 20 µM UbVS, 5 µM NEL domain (E3) at RT for 2 hours.

However, we also observed differences between the activities of these proteins. Labelling with UbVS was different between the four NEL family members: both FL and ΔN constructs of IpaH9.8 and IpaH1.4 were approximately 50% labelled after two hours, whereas SspH2 was almost completely labelled, in contrast to SspH1 which showed the lowest level of labelling (Figure 1F, Supplementary Figure 2D). With the NEL domains alone, labelling was more similar across the family (Figure 2C), though SspH1 still showed the lowest degree of labelling. In NEL domain-only ubiquitination assays, we observed a similar propensity to form free ubiquitin chains with IpaH1.4 and IpaH9.8, while SspH1 and SspH2 appeared less active (Figure 2A).

Similarly, time course E2∼Ub discharge assays with ΔN constructs highlighted that IpaH1.4 and IpaH9.8 were much more efficient at discharging ubiquitin than SspH1 or SspH2 (Figure 1D and E). To observe complete ubiquitin discharge within the same time frame, a 10-fold higher concentration of SspH proteins was required when compared to the IpaH proteins. This corroborates previous work by Keszei and Sicheri, where longer assay time points were used to study SspH1 activity compared to IpaH9.8.^30^ For E2∼Ub discharge assays with isolated NEL domains, we observed similar rates of free di-ubiquitin synthesis with IpaH1.4 and IpaH9.8, while SspH2 was significantly less efficient compared to the other three proteins (Figure 2B). Interestingly, the E3∼Ub thioester intermediate was sufficiently stable in these assays that it could be observed at early time points with IpaH1.4, IpaH9.8 and SspH1.

We found it intriguing that SspH1 is the apparently least accessible of the NELs tested to ubiquitin-loading with UbVS (Figure 1F) and has a lower turnover in discharge assays compared to IpaH proteins (Figure 1D and E, Figure 2B) but is the most active in substrate ubiquitination assays (Figure 1B). Therefore, we tested whether the presence of PKN1 HR1b substrate might alter the dynamics of SspH1 and therefore accessibility of the catalytic cysteine. However, we observed that PKN1 had no effect on SspH1 loading with UbVS ubiquitin (Supplementary Figure 4A). This is complementary to, and consistent with E2∼Ub discharge assays performed by Keszei and Sicheri where the addition of PKN1 HR1b to SspH1 had no impact on E2∼Ub discharge rate, indicating that substrate binding does not significantly alter access to the SspH1 active site.^30^

### ΔNSspH1 crystal structure

To better understand the mechanism of SspH1 and the apparent differences in activity, we solved the crystal structure of a construct containing the LRR and NEL domains but lacking the first 160 residues, ΔNSspH1 (now referred to as SspH1). The structure was refined at 2.9 Å resolution with one copy in the asymmetric unit and a *P*622 space group. Details of data collection and statistics are reported in Supplementary Table 1.

The structure is elongated with the LRR (residues 161-393) domain extending away from the NEL domain (residues 405-622) which contains the catalytic cysteine (C492) and the α-helices of the E2-Ub binding domain (E2-UbBD, residues 623-697) (Figure 3A). It is similar to IpaH9.8 and IpaH3 in an ‘open’ conformation, with the position of the E2-UbBD relative to the catalytic loop in an analogous position to other bacterial E3 ligase structures (Figure 3B). The E2-UbBD domain of SspH1 consists of two long anti-parallel α-helices and a small C-terminal helical segment as also seen in SspH2 (Figure 3C and Supplementary Figure 1A). In contrast, IpaH9.8, IpaH1.4 and SlrP have a notable kink in one of the two long anti-parallel α-helices (Figure 3C). In agreement with our sequence alignment (Supplementary Figure 1A), SspH1 lacks the amplifier loop in LRR6 of the IpaH9.8 LRR domain which was shown to be important for GBP1 substrate sensing.^29^ SspH2 also lacks an amplifier loop, while IpaH3 is missing electron density for eight amino acids in the corresponding LRR, and SlrP has an extended loop in the next LRR.^9,28^ Whether these loops act as signaling amplifiers upon substrate binding is unknown.

**Figure 3.**
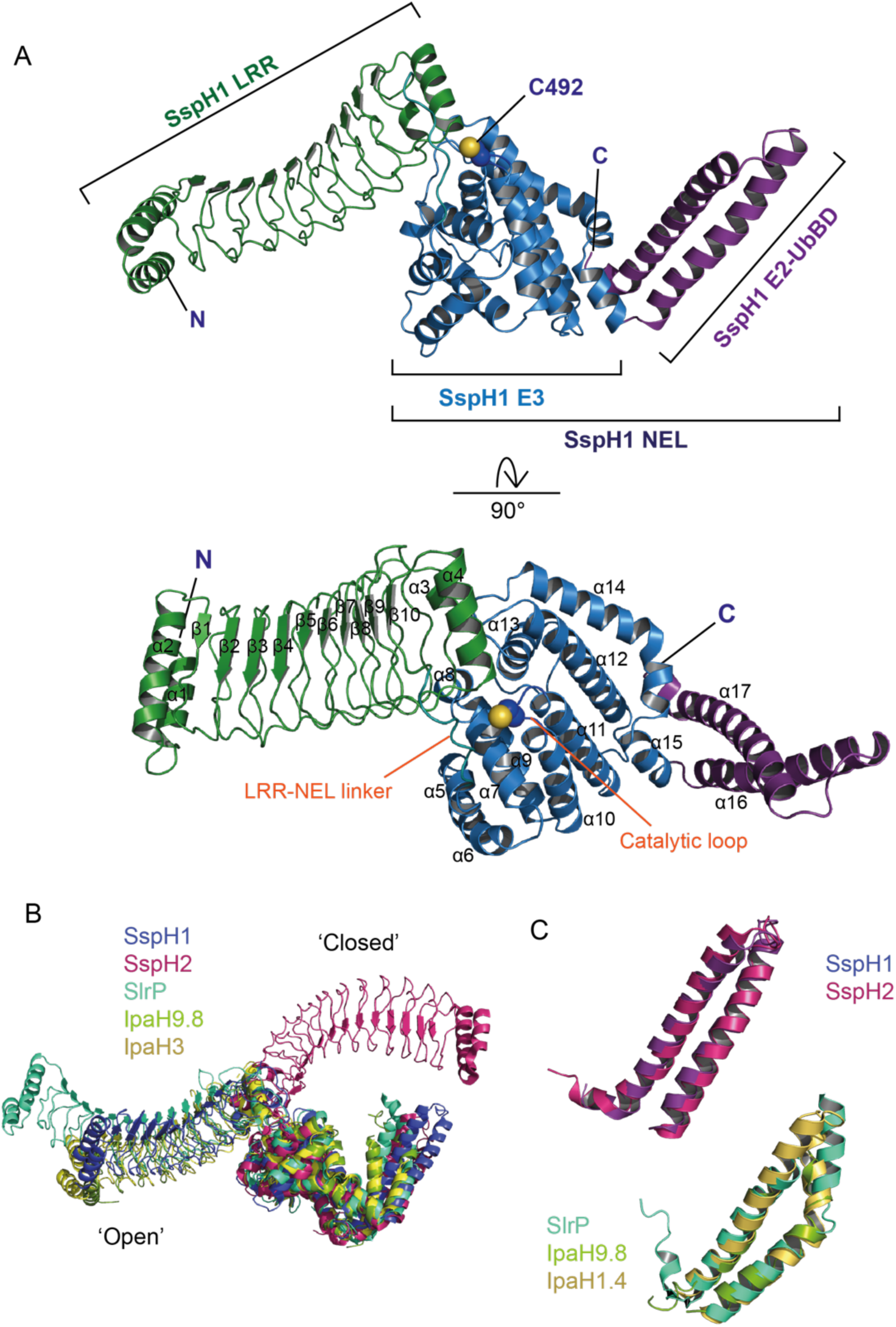
Crystal structure of ΔNSspH1. A) Overall structure from side and top view; B) Loop containing catalytic cysteine, with the Cysteine side chain shown as spheres; C) Overlay of ΔN NEL structures aligned to the NEL domain showing SspH1, SlrP (PDB 3PUF), IpaH9.8 (PDB 6LOL) and IpaH3 (PDB 3CVR) in the ‘open’ conformation, and SspH2 (PDB 3G06) in the ‘closed’ conformation; D) Aligned E2-UbBD thumbs for SSpH1 and SspH2, and SlrP, IpaH9.8 and IpaH1.4 (PDB 3CKD).

In our SspH1 structure, sixty N-terminal residues, those in the LRR-NEL linker (394-404) and the loop containing the catalytic C492 have a higher B-factor compared to other residues suggesting more flexibility in these regions. A highly dynamic LRR-NEL linker and catalytic loop might be an important mechanistic feature in bacterial E3 ligases as high B-factors or lack of electron density are also observed for these regions in other available NEL structures that contain both LRR and NEL catalytic domains: ΔN SspH2, IpaH3 and IpaH9.8 do not have electron density for the LRR-NEL linker, while IpaH9.8 is also missing density for the catalytic loop.^9,28,29^ SlrP, is currently the only available structure of an LRR-NEL construct that has its substrate thioredoxin bound. The complex crystallized as a heterotetramer, with the two SlrP molecules in an open, head to tail arrangement that are bridged by 2 molecules of thioredoxin-1, though it is not known if self-association is required for substrate ubiquitination by SlrP. In this structure there is well-defined density for the loop linking the LRR and NEL domains which makes contacts with Trx-1.^35^

We observed a symmetry related molecule in our structure of SspH1 where the E2-UbBD of one molecule sits on the top of the LRR domain, close to its substrate binding interface (Supplementary Figure 5A). To determine if this was a biologically relevant interaction or an artefact of crystal packing, we performed SEC (size-exclusion chromatography)-MALLS (multi-angle laser light scattering measurements) with FL SspH1 and ΔNSspH1. We confirmed that, in the range of concentrations explored, both are monomeric in solution (Supplementary Figure 5B and C), and form a 1:1 complex with the substrate PKN1 (Supplementary Figure 5D and E), in agreement with a previous report.^11^ This is in contrast to SlrP which has been shown to exist in a monomer-dimer equilibrium.^35^

We next interrogated the importance of residues in SspH1 that make contacts between the LRR and NEL domains to better understand the interplay between these two domains and their role in autoinhibition. R351, which is conserved in SspH1/2 and SlrP but not IpaH proteins forms key interactions with E493 and D552, while R375, which is an Arg or His across the NEL family (Supplementary Figure 1) points towards an acidic pocket formed by residues surrounding the catalytic acid and base (Figure 4A and Supplementary Figure 6A).^30^ However we observed no significant difference between mutants R351A, R375A, R351A R375A and WT SspH1 activity in either auto-ubiquitination or PKN1 ubiquitination assays (Figure 4B), indicating that these residues do not regulate autoinhibition. Additionally, hydrophobic interactions with L550 at its center further contribute to the LRR-NEL interface (Supplementary Figure 6A). This observation supports previous mutational analysis of L550 which showed that L550S partially released autoinhibition but had no apparent effect on substrate ubiquitination ^27,36^. Interestingly, the L550D mutation changed the pattern of substrate ubiquitination, suggesting that this position affects the conformational changes required to bring the substrate close to the E3∼Ub thioester (Supplementary Figure 6B).

**Figure 4.**
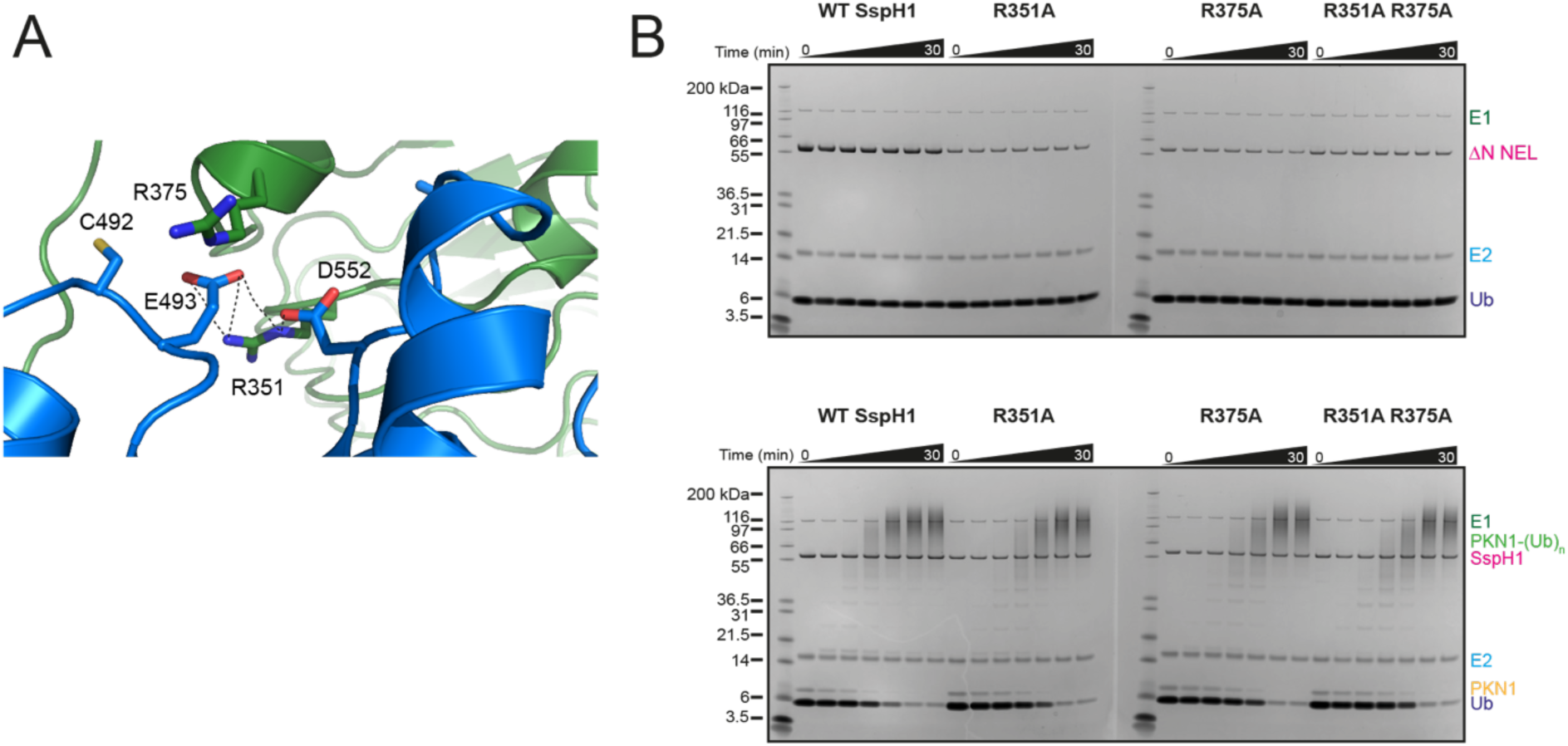
Interface between the LRR and NEL domains. A) Interface between the LRR domain in green and the NEL domain in blue highlighting the interactions made between R351 (LRR) and E493 and D522 (NEL). The side chain of the catalytic cysteine is shown. B) Top: Auto ubiquitination assay with ΔNSspH1 mutants R351A, R375A, R351A R375A. Performed with 0.1 µM UBA1 (E1), 2 µM UbcH5A (E2), 1.0 µM SspH1 (E3), 20 µM ubiquitin, 10 mM ATP at RT for 0-30 minutes. Bottom: Substrate ubiquitination assay with ΔNSspH1 mutants R351A, R375A, R351A R375A. Performed with 0.1 µM UBA1 (E1), 2 µM UbcH5A (E2), 1.0 µM SspH1 (E3), 2 µM HR1b PKN1, 20 µM ubiquitin, 10 mM ATP at RT for 0-30 minutes.

At present, the SlrP/Trx-1 complex is the only available structure of a near full-length NEL bound to its substrate. However, the tetrameric nature of this complex makes it difficult to draw general conclusions about the mechanism of substrate ubiquitination, given that other NEL family members are assumed to transfer ubiquitin in a 1:1 NEL/substrate complex. Furthermore, no structure of apo SlrP is available to assess conformational changes induced upon substrate binding. Instead, near full-length IpaH9.8 has been structurally characterized in the apo state and the isolated LRR domain bound to its substrate GBP1 providing first insight into the effect of substrate engagement.^29^ Comparison of these two structures allowed identification of structural changes induced in the LRR upon substrate binding, especially a rotation of the LRR-CT toward the convex side of the LRR, which destabilizes a hydrophobic cluster in the C-terminus of the LRR that engages F395 (equivalent to L550 in SspH1) from the NEL, thereby enabling release of the NEL domain and autoinhibition. The structure of SspH1 presented here, now allows us to carry out the same comparison using the previously published structures of the SspH1 LRR in its apo form and in complex with its substrate PKN1 HR1b.^27^ Interestingly, an overlap of the PKN1-bound LRR with SspH1 revealed that the changes induced upon substrate binding are much smaller than those observed in IpaH9.8 with an RMSD of 0.685 Å (Supplementary Figure 6C), without any significant effect on the hydrophobic environment of L550. This suggests that substrate binding to SspH1 alone might not be sufficient to induce the conformational changes required to release autoinhibition.

### Solution properties of FL and ΔNSspH1, and substrate-bound complexes

The observation that LRR and NEL domains adopt different orientations with respect to one another in crystal structures of apo NEL proteins indicates the presence of significant interdomain flexibility, however little is known about substrate binding affects this conformational flexibility. To investigate the solution properties of SspH1 we recorded Small-angle X-ray Scattering (SAXS) data on full-length (FL) and ΔNSspH1 constructs in the presence and absence of PKN1 (Figure 5A-B and Supplementary Figure 7). Details of data collection and analysis are reported in Supplementary Table 2.

**Figure 5.**
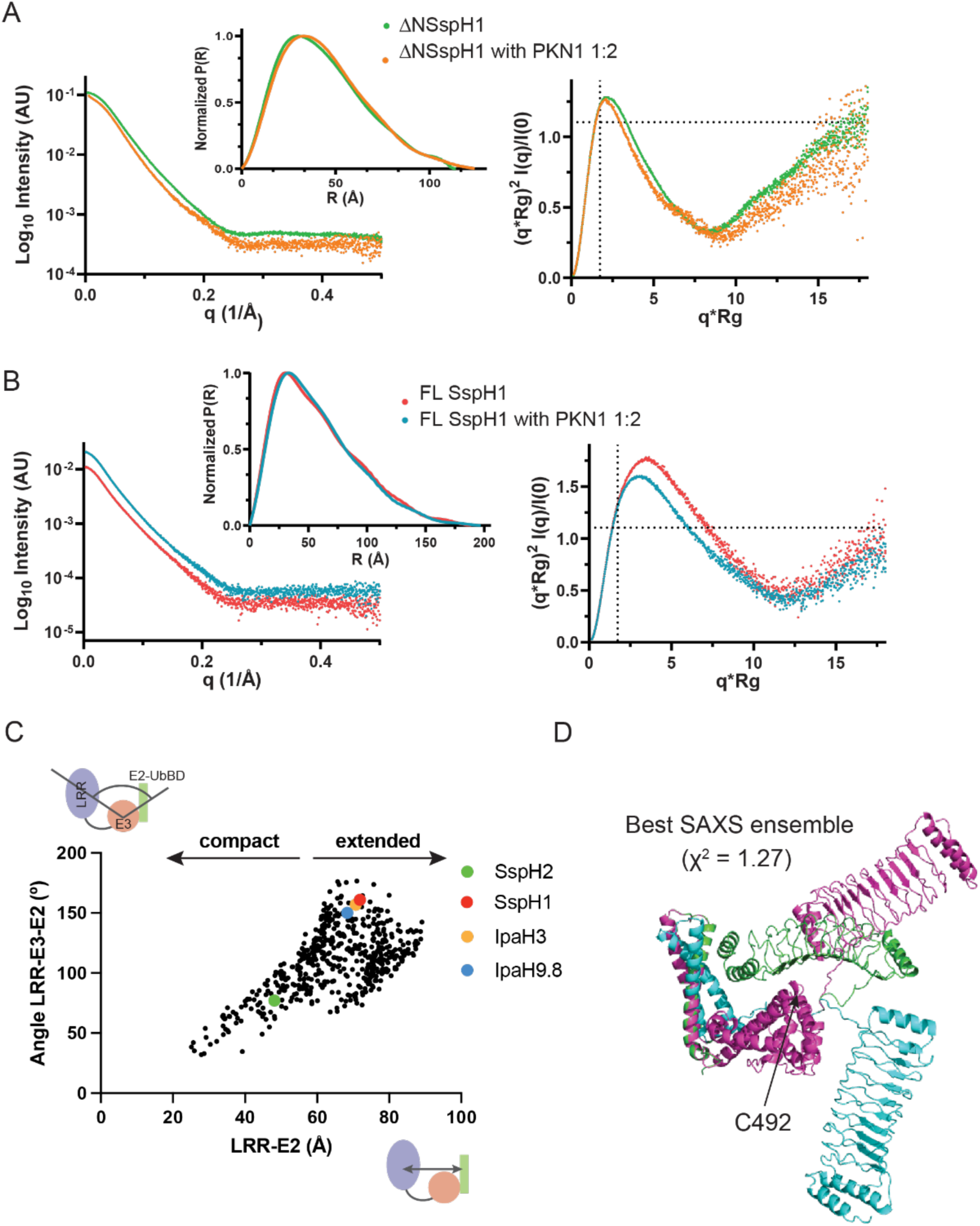
Small-Angle X-ray scattering analysis. X-ray scattering intensities profiles, normalized pair distance distribution P(R) and Kratky plot of ΔSspH1 (A) of full-length SspH1 (B) in the absence and presence of PKN1. The dotted lines in the Kratky plot are drawn at qR_g_ = ✓3 and (qR_g_)^2^I(q)/I(0) = 1.104. Folded and globular proteins have a maximum where the two lines intersect.^50^ (C) LRR-NEL intra-domain angles and distances distribution for the structural ensembles calculated by Xplor-NIH that represent the experimental Xray scattering intensities (χ2 ≤ 1.5). (D) The three-conformers structural ensemble that best agrees with the experimental SAXS data (lowest χ^2^).

We observed a 75 Å decrease in the maximum dimension (Dmax) and a 14 Å reduction in the radius of gyration (Rg) for ΔNSspH1 compared to the FL protein indicating a large contraction in the protein volume following the deletion of the small N-terminal domain. Interestingly, the change in molecular dimensions is not associated with a change in the respective radii of cross-section (Rc) suggesting that the overall structures maintain elongated shapes (Dmax>>Rg) of similar widths. Rather surprisingly, the Guinier analysis of ΔNSspH1 and FL protein scattering data reveals similar radii of gyration for both proteins in isolation and bound to the HR1b domain of PKN1 (Supplementary Figure 7A). A 10 Å increase in the maximum dimension of the normalized P(R) distributions is observed for both constructs following substrate recruitment, suggesting that only small molecular rearrangements occur upon PKN1 binding. In both cases the molecular weights calculated from the SAXS data are those expected from the primary sequences suggesting the proteins in their apo forms and in complex with PKN1 are monomeric in solution with a 1:1 stoichiometry, as reported previously and in agreement with our SEC-MALLS analysis (Supplementary Figure 5).^11^

We assessed the flexibility of the protein constructs under investigation using dimensionless Kratky plots (Figures 5A and B). The broad bell shape with a maximum shifted towards higher q*Rg values, especially apparent in FL SspH1, together with the long tails in the P(r) distributions, confirms that SspH1 populates elongated conformations. Moreover, the uptrend of the Kratky profile at higher q*Rg indicates the presence of overall conformational flexibility which is unaffected by the presence of the substrate. Fitting the solution scattering data using either our ΔNSspH1 crystal structure coordinates or an AlphaFold3 model, both of which are characterized by distinct orientations of the LRR and E2-UbBD relative to the catalytic E3 domain, yields large χ2 values in both cases. This indicates that neither structure accurately represents the solution conformation of SspH1, most likely due to its dynamic features. To explore the conformational space available to ΔNSspH1 we modelled solution structure ensembles based on the experimental X-ray scattering data using an Xplor-NIH implemented protocol.^37^ Ensembles were chosen based on the agreement with the experimental data (χ2 ≤ 1.5) and visualized in a plot reporting the angles between the centers of mass of the E2-UbBD, E3 and LRR domains as a function of the distances between the E2-UbBD and LRR centers. Relative positions of the same domains in our ΔNSspH1 crystal structure, as well as the available structures of ΔNSspH2, IpaH3 and IpaH9.8 are also reported. ΔNSspH1 ensembles conformers possess significant interdomain flexibility, implying that the protein can adopt both extended (open) and more compact (closed) conformations (Figure 5C). Structure envelope maximum dimensions (Supplementary Figure 7C) and radii of gyration (Supplementary Figure 7D) for the crystal structures of ΔNSspH1, ΔNSspH2, ΔNIpaH9.8 and ΔNIpaH3 all fall within overall ensemble distributions for these parameters. Notably, the ensemble best agreeing with the experimental scattering data (lowest χ2 value) comprises two extended structures and a closed conformation, wherein the substrate binding interface on the LRR domain is oriented toward the catalytic loop (Figure 5D).

In conclusion, our SAXS data reveal that SspH1 exhibits significant interdomain flexibility suggesting that available crystal structures likely represent specific conformational snapshots within a continuous structural landscape. While PKN1 binding does not fundamentally alter SspH1 overall flexibility, we observed a subtle shift in the Kratky profile maximum towards smaller q*Rg for both the FL protein and ΔNSspH1 upon PKN1 binding (Figures 5A and B). This suggests that PKN1 binding might modulate the distribution of SspH1 closed and open conformations resulting in a small increase in globularity of the average solution structure of the complex compared to the apo states.

### Dynamic properties of the SspH1-ubiquitin conjugate

We sought to better understand the structural flexibility of ubiquitin when loaded to SspH1 alone (SspH1∼Ub) and in complex with PKN1. Our SAXS analysis indicated subtle changes in SspH1 conformation upon PKN1 binding. Given the low efficiency of UbVS labelling (Figure 1F), which we used to form a stable SspH1-Ub covalent conjugate, we employed NMR to study how E3 conjugation affects ubiquitin dynamic behaviour.

We first examined the non-covalent interaction of SspH1 with ^15^N-labelled ubiquitin. by monitoring the effect of SspH1 addition on the backbone amide resonances in the ^1^H-^15^N HSQC spectrum of an ^15^N-labelled ubiquitin sample (Figure 6). Fast exchange in a subset of ubiquitin backbone amide resonances was observed upon SspH1 addition (Figure 6A). The perturbed resonances are located at the ubiquitin N- and C-terminal residues and amide protons around the ubiquitin I44 hydrophobic patch (Figure 6D and G). Incremental line broadening was also observed for the perturbed resonances in the ubiquitin spectrum, likely due to the increase in molecular size and longer correlation times upon interaction with SspH1. Next, we probed the interaction of the ubiquitin with the individual SspH1 domains. Chemical shift perturbations (CSPs) were still visible but fewer resonances were affected in the ^1^H-^15^N HSQC spectrum when the isolated NEL domain was added to ^15^N-labelled ubiquitin compared to SspH1 (Figure 6B). The differences in the two interfaces are located mainly in the ubiquitin C-terminal tail (Figure 6E and H). Resonance line broadening is also less pronounced in the titration of ^15^N-ubiquitin with the NEL alone compared to SspH1 (Figure 6A and B). These observations point to a direct participation of the LRR domain in the SspH1/ubiquitin interaction. In fact, a titration of ubiquitin with SspH1 featuring a largely flexible and independent LRR domain would have shown the same profile of CSPs and line broadening observed for the titration of ubiquitin with the isolate NEL domain. Addition of the LRR domain alone to ^15^N-labelled ubiquitin led to a largely unperturbed ^1^H-^15^N HSQC spectrum, with some cross peaks experiencing small CSPs and associate resonance broadening (Figure 6C). A residue-specific line-broadening analysis identified residues primarily located in the N- and C-terminal regions of ubiquitin experiencing larger effects upon interaction with the LRR (Figure 6F and I). While this LRR/Ub interaction is weak, its surface appears to be complementary to the NEL/Ub interface, suggesting the two domains use different ubiquitin surfaces for their interaction.

**Figure 6.**
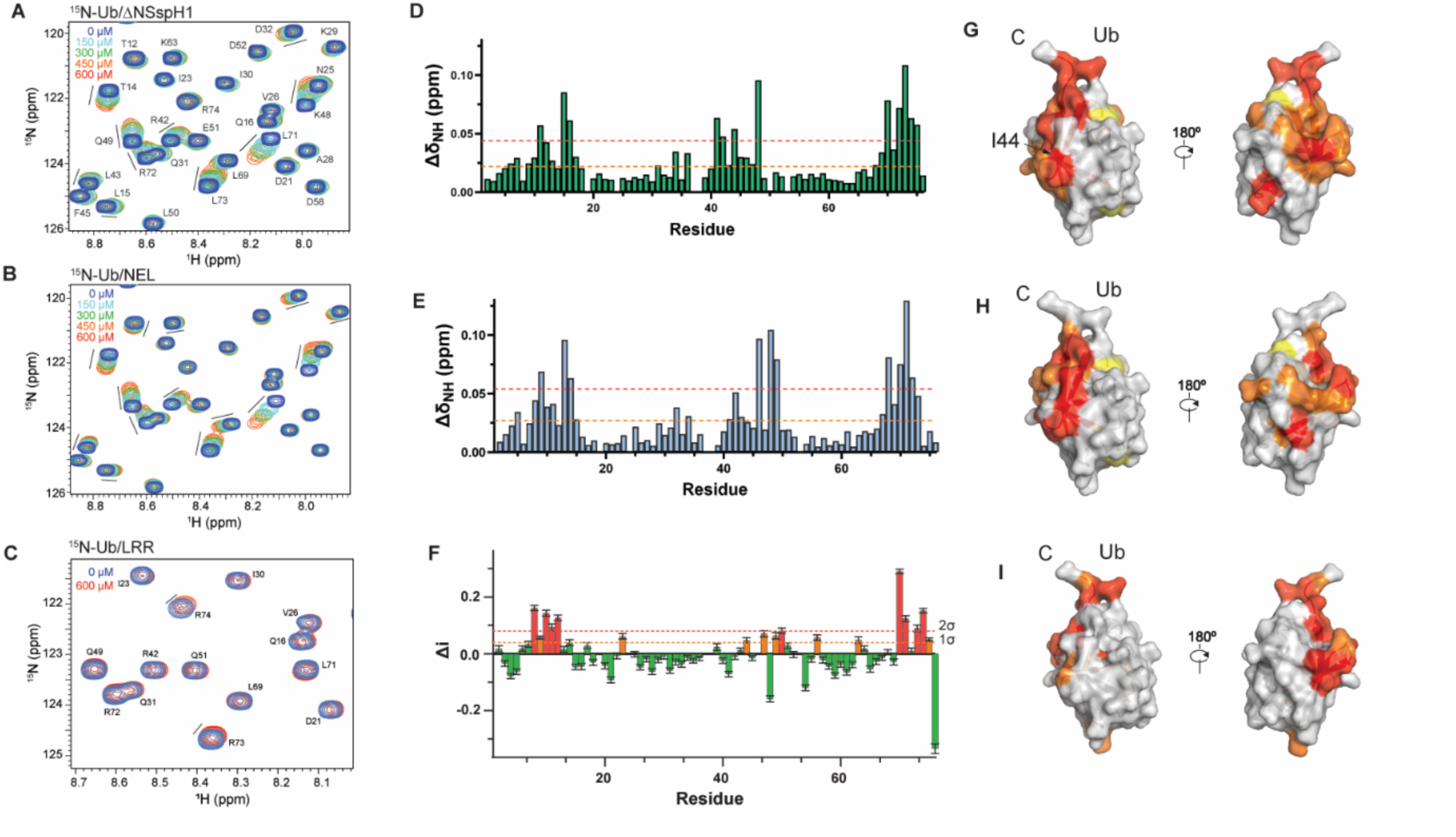
NMR analysis of the SspH1-ubiquitin interaction. Details of the titration of ^15^N-labeled ubiquitin titrated with ΔN SspH1 (A) SspH1 NEL domain (B), and SspH1 LRR domain (C). Spectra at different ligand concentrations are plotted at the same contour level. Chemical shifts perturbations versus residue number in the NMR spectrum of ^15^N-labeled ubiquitin in the presence of (D) ΔN SspH1 or (E) NEL domain with CSPs mapped onto ubiquitin structure (G & H). Ubiquitin ^1^H-^15^N HSQC line broadening versus residue number for the titration with the LRR domain with the rate of broadening mapped onto ubiquitin structure (F&I).

To try and rationalize these observations we used AlphaFold3 (AF3) to model the structure of ΔNSspH1 bound to Ub (Supplementary Figure 8A).^38^ In the predicted SspH1/Ub complex, the SspH1/Ub interface aligns with the cluster of ubiquitin residues experiencing CSPs observed in the NMR titration experiments with the ΔNSspH1 and NEL domain. To investigate whether SspH1 uses the predicted surface to interact with ubiquitin, we mutated a glutamate residue (E487) in SspH1 that appears to serve as a linchpin for binding, being packed between the side chains of R72 and R42 of ubiquitin (Supplementary Figure 8A). We hypothesized that, if this was the correct E3/Ub binding mode, E487 mutation to alanine would weaken ubiquitin binding, while mutation to arginine would completely abrogate it. Whilst there was no difference in the effect on the backbone amide resonances in the ^1^H-^15^N HSQC spectrum of a ^15^N-labelled ubiquitin sample upon addition of either WT or E487R SspH1 (Supplementary Figure 8B), we observed disrupted discharge and substrate ubiquitination activity for the E487A mutant, and its ablation for the E487R mutant (Supplementary Figure 8C and D). This indicates that whilst E487 has a role in SspH1 activity, the AF3 predicted interface between the ubiquitin and SspH1 is likely not the primary function of this residue. Instead, E487 interacts with H498 which has previously been shown to be important for the transfer of Ub from E3∼Ub to the substrate (Supplementary Figure 8E).^11^ Interestingly, these two residues are located on the helices either side of the catalytic loop containing C492 suggesting that their interaction may stabilize its position to support catalysis.

Next, we investigated the dynamic behavior of ubiquitin in the SspH1∼Ub conjugate and how it is altered by PKN1 binding. We recorded ^1^H-^15^N HSQC spectra of SspH1 pre-loaded with ^15^N-labelled ubiquitin UbVS probe and monitored the effect on ubiquitin resonances in the SspH1∼Ub conjugate when PKN1 is added (Figure 7). While the charging of SspH1 with ^15^N-ubiquitin probe is inefficient, this approach is suitable for NMR experiments as only the charged ^15^N-labelled ubiquitin in the SspH1∼Ub conjugate is visible in the spectra. We first confirmed that the ^1^H-^15^N HSQC spectrum of the free ^15^N-ubiquitin UbVS probe remains largely indistinguishable from that of the wild-type protein, except for a few C-terminal residues, implying that the addition of the vinyl sulfone warhead does not induce structural changes in ubiquitin. Upon conjugation to the SspH1 catalytic cysteine, most backbone resonances in the ^15^N-labeled ubiquitin ^1^H-^15^N HSQC spectrum probe decrease in intensity with a subset experiencing a stronger line broadening effect (Figure 7A). The remaining visible cross peaks (backbone and side chain amide resonances) in the ^1^H-^15^N-HSQC spectrum show no change in their chemical shifts. A possible interpretation of this observation is that in the SspH1∼Ub conjugate, the ubiquitin exists in fast exchange between two states: a predominant, flexibly tethered state, visible by NMR, and a less abundant, locked state, invisible in the ^1^H-^15^N HSQC ubiquitin spectrum due to its high molecular weight (∼70 kDa). This equilibrium would explain the observed pattern of line broadening and the persistence of some visible peaks. It suggests that while ubiquitin is not permanently locked in a rigid conformation with SspH1, it does transiently become locked in a larger complex, possibly involving interactions with both the NEL and LRR domains. Upon addition of PKN1 substrate to the SspH1∼Ub conjugate, there is an overall increase in the ubiquitin cross peaks intensity and a subset of ubiquitin resonances, that were previously invisible or weak, either reappear or increase in intensity (Figure 7B). This indicates a shift in the fast exchange equilibrium between the locked and flexible ubiquitin states towards a more dynamic ubiquitin. This shift may result from the loss of one or more interaction surfaces between ubiquitin and SspH1 upon PKN1 binding. Intriguingly, the resonances that showed increased intensity cluster to a patch on the ubiquitin molecule, encompassing the I44 hydrophobic surface, that overlaps with the region found to interact with SspH1 in our NMR results with free ubiquitin (Figure 6G and H).

**Figure 7.**
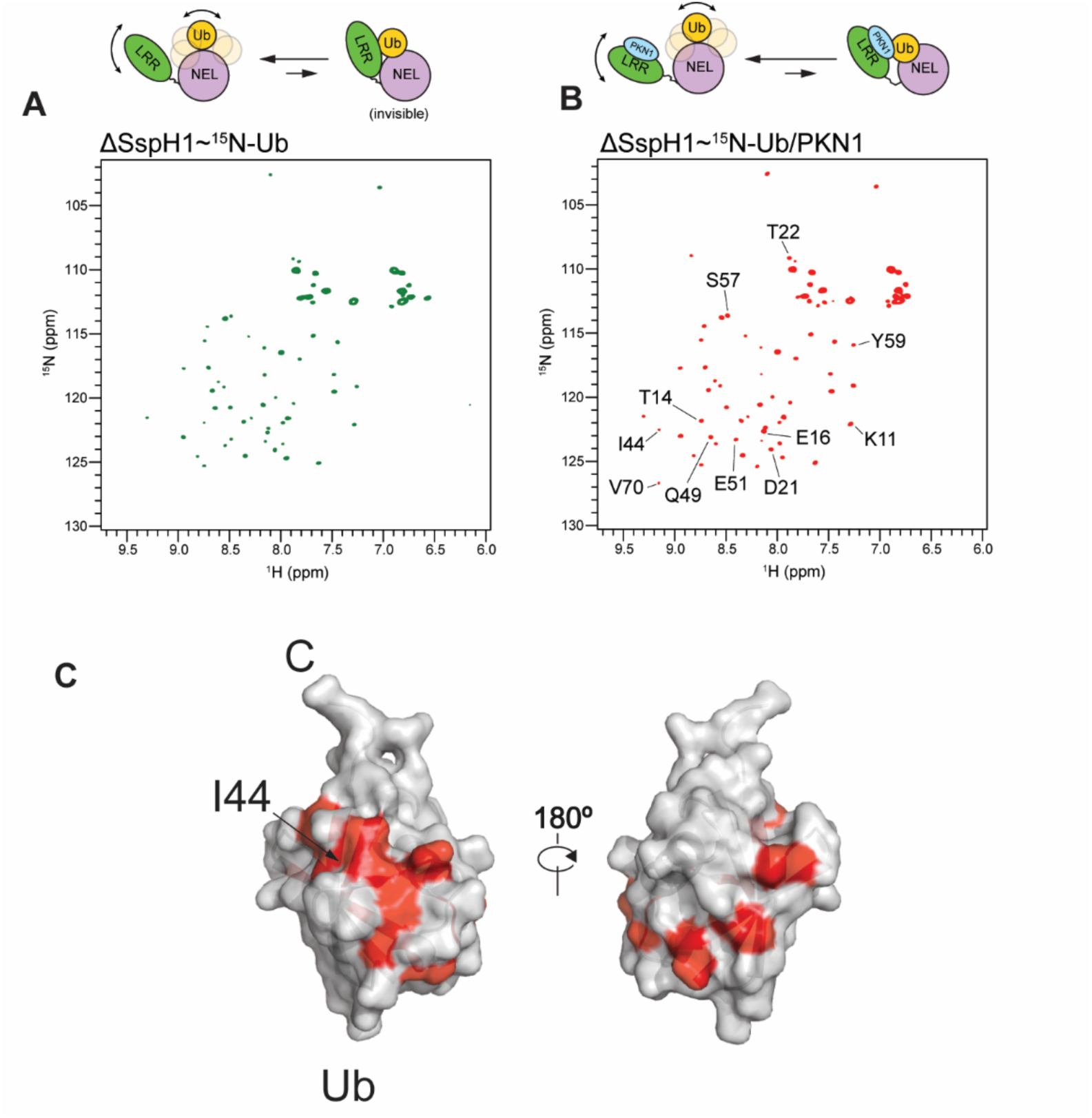
NMR of conjugated^15^N-labelled ubiquitin. Spectra of ^15^N-Ub tethered to ΔNSspH1 C492 in the absence (A) and presence (B) of PKN1. Residues in the ubiquitin ^1^H-^15^N HSQC spectrum reappear upon addition of the substrate. The reappearing residues are plotted on the ubiquitin structure in (C).

Taken together, our NMR experiments show that ubiquitin non-covalently interacts with both the NEL and LRR domains of SspH1. Our results with tethered ubiquitin, which mimics the activated E3∼Ub intermediate, suggest that, in the SspH1∼Ub conjugate, the LRR domain restricts ubiquitin dynamic behaviour and substrate binding impacts ubiquitin flexibility in the SspH1∼Ub/PKN1 complex.

## DISCUSSION

Bacterial NEL E3 ligases have an important role in *Salmonella* and *Shigella* evasion of host cell immune responses during infection, however the exact molecular mechanism of ubiquitin transfer and regulation of activity remains unclear. In this study, we performed a cross-species comparison of NEL catalytic activity. We show that active site cysteines of full length IpaH and SspH proteins are readily accessible for ubiquitin loading but are immediately discharged, confirming that autoinhibition is not due to steric blockage of the catalytic cysteine, and supporting the model that water is the favored nucleophile for ubiquitin discharge in the absence of substrate.^30^ This activity is advantageous to the bacteria, since it prevents auto-ubiquitination and proteasomal degradation of the NELs themselves, and unanchored ubiquitin chain formation, which could trigger the host cell innate immune response. However, we also identified differences in activity between *Salmonella* and *Shigella* NELs. To better understand these differences, we solved the crystal structure of a SspH1 construct containing both LRR and NEL domains. When compared to previously published structures for SspH2, IpaH9.8 and IpaH3, SspH1 is most similar in orientation to IpaH9.8 and IpaH3, with an ‘open’ conformation.

Previous work had shown that NEL family members rely on significant flexibility within the NEL domain to recruit ubiquitin.^11^ While available crystal structures of NEL proteins and substrate complexes provide snapshots of specific states, little is known about how ubiquitin loading affects conformational dynamics of the ligase, how substrate binding affects the overall protein dynamics of NELs and how ubiquitin is passed onto the substrate. Here, we have characterized the solution state behaviour of SspH1 by SAXS and show that it exists in a conformational continuum, which include the orientations captured in the crystal structures of SspH2, IpaH9.8 and IpaH3. Furthermore, our SAXS analysis shows that apo SspH1 has a significant level of flexibility around the hinge between the LRR and NEL domains, which is not substantially affected by substrate binding, though it shifts the equilibrium towards more globular conformations of SspH1. NMR experiments with free ^15^N-labelled ubiquitin and a ^15^N-labelled ubiquitin probe loaded onto the catalytic cysteine of SspH1 enabled us to study the environment of ubiquitin in the presence of SspH1 and PKN1. These experiments suggest that ubiquitin is partially restrained when non-covalently or covalently bound to SspH1, forming transient interactions with the NEL domain and possibly weaker interactions with the LRR domain. The interactions with the LRR domain are relieved upon PKN1 binding.

The crystal structure of the SspH1 construct containing both the LRR and NEL domains allowed comparison with the previously reported substrate-bound LRR domain structure, to identify potential changes induced upon substrate binding, as previously described for IpaH9.8. Interestingly, no significant structural changes are induced by the substrate, indicating that the mechanism of release of autoinhibition suggested for IpaH9.8 must be protein specific and not generally applicable to this protein family. Instead, we identified E487 in the NEL as important for activity. This residue interacts with H498 from the adjacent helix which has previously been reported to be important for ubiquitin transfer from E3 onto the substrate and we speculate that interaction between these two residues might be important for stabilizing the catalytic loop.

Taken together with previously published studies, the work presented here suggests that substrate binding to the LRR domain of NEL proteins is not sufficient to induce the major conformational changes required to bring the substrate close to the E3∼Ub thioester intermediate to enable ubiquitin transfer. At present it is unknown what additional events are necessary to promote formation of a ubiquitin transfer competent conformation of a substrate-NEL∼Ub complex and further studies are required to trap members of this enigmatic E3 ligase family in their active state.

## Acknowledgements

The authors would like to thank Andrew Purkiss for crystallography data collection, Aurelien Thureau for SAXS beamline support, and Ian Taylor for help with SEC-MALLS experiments. X-ray crystallography data collection was performed on the i24 beamline at Diamond; SAXS data collection on the SWING beamline at Soleil synchrotrons. This work was supported by the Francis Crick Institute, which receives its core funding from Cancer Research United Kingdom (CC 2075), the United Kingdom Medical Research Council (CC 2075), and the Wellcome Trust (CC 2075); by the Biotechnology and Biological Research Council, BB/T014547/1 to K.R. and D.H.; and by the Engineering and Physical Sciences Research Council, EP/V038028/1 to D.H. and K.R. For the purpose of open access, the author has applied a CC BY public copyright license to any author-accepted manuscript version arising from this submission. Figures were created in BioRender.com under the institutional license belonging to the Francis Crick Institute.

## Author contributions

C.R.K and D.E.: experiments, data analysis, visualisation, writing original draft, review & editing. D.H. and K.R.: conceptualisation, data analysis, funding acquisition, supervision, writing – review & editing.

## Methods

### Protein Expression and Purification

All proteins were expressed in pET49b vectors containing a 3C cleavage site and His-tag in BL21 Gold *E.Coli* in LB media. Cells were grown at 37 C until OD600 reached 0.6-0.8, and protein expression induced with 0.5 mM IPTG overnight at 20 C. Cells were lysed by sonication in 50 mM HEPES pH 7.5, 150 mM NaCl, 20 mM imidazole, 0.5 mM TCEP in the presence of protease inhibitor. Proteins were purified on nickel NTA resin (Thermo Scientific, #88222), followed by elution and treatment with 3C protease overnight. Proteins were then purified by gel filtration into 50 mM HEPES pH 7.5, 150 mM NaCl, 0.5 mM TCEP. The following constructs were used in activity assays: Full length constructs: IpaH1.4 1-575, IpaH9.8 1-545, SspH1 1-700, SspH2 1-788. ΔN constructs: IpaH1.4 31-575, IpaH9.8 21-545, SspH1 161-700, SspH2 166-788. NEL domain constructs: IpaH1.4 283-575, IpaH9.8 252-545, SspH1 405-700, SspH2 493-788.

### Substrate ubiquitination assays

NEL E3 ligase constructs of IpaH1.4, IpaH9.8 and SspH1 (1 µM) were incubated at 22 degrees with shaking with their respective substrates LUBAC (C82A, 1 µM), GBP1 (1 µM) and PKN1 (residues 122-199, 2 µM) in the presence of 0.1 µM E1, 2 µM UbcH5A, 20 µM ubiquitin and 10 mM ATP in 25 mM HEPES pH 7.5, 150 mM NaCl, 10 mM MgCl_2_ for 0-30 minutes. Samples were quenched with LDS loading dye containing DTT and run by SDS-PAGE on 4-12% gels.

### Auto-ubiquitination Assays

NEL E3 ligase constructs of IpaH1.4, IpaH9.8, SspH1 and SspH2 (1 µM) were incubated at 22 degrees with shaking in the presence of 0.1 µM E1, 2 µM UbcH5A, 20 µM ubiquitin and 10 mM ATP in 25 mM HEPES pH 7.5, 150 mM NaCl, 10 mM MgCl2 for 0-30 minutes. Samples were quenched with LDS loading dye containing DTT and run by SDS-PAGE on 4-12% gels.

### Discharge Assays

NEL E3 ligase constructs of IpaH1.4, IpaH9.8, SspH1 and SspH2 (100 nM) were incubated at 22 degrees with shaking in the presence of 1 µM precharged UbcH5A-Ub-cy3 for 0-20 minutes. Samples were quenched with LDS loading dye and run by SDS-PAGE on 4-12% gels.

### UbVS loading assays

NEL E3 ligase constructs of IpaH1.4, IpaH9.8, SspH1 and SspH2 (5 µM) were incubated with 20 µM ubiquitin vinyl sulfone (UbVS, U-202B-050, BioTechne) at 22 degrees with shaking for 0-120 minutes. Samples were quenched with LDS loading dye containing DTT and run by SDS-PAGE on 4-12% gels.

### Crystallization, data collection, phasing and refinement

ΔNSspH1 (161-700) at a concentration of 12 mg/mL was crystallized in 50 mM HEPES pH 7.5, 150 mM NaCl and 0.5 mM TCEP at 20 °C. Crystals grew from a sitting drop made of 0.1 µl of protein solution mixed with 0.1 µl of 0.225 M sodium tartrate, 22% PEG 3350, 11 mM sarcosine. The crystals were cryo-protected with 0.225M sodium tartrate, 35% PEG 3350, 10% glycerol and data were collected at Diamond Light Source at the i24 beamline. Data processing was done using DIALS and merged and scaled using AIMLESS.^39,40^ The initial model was obtained by molecular replacement using the available coordinates of the SspH1 LRR domain (4NKH) and the coordinates of the NEL domain from the SspH1 AlphaFold model as templates in Phenix Phaser.^41^ Models were iteratively improved by manual building in Coot and refined using both REFMAC and Phenix.^41–43^ Coordinates and structure factors are deposited in the PDB with the code 9H6W and the final refinement statistics are reported in Supplementary Table 1.

### Small-Angle Xray Scattering and SspH1 ensemble modelling

SAXS data were collected at the SWING beamline at SOLEIL (GIF-sur-YVETTE CEDEX, France). Samples of ΔNSspH1 and FL SspH1 at concentrations 168 µM and 104 µM in 50 mM HEPES pH 7.5, 150 mM NaCl and 0.5 mM TCEP in the absence and presence of 1:2 molar equivalents of PKN1, were injected onto a Bio SEC-3 100 Å Agilent column and eluted at a flow rate of 0.3 ml/min at 15 °C (Figure S8A). Frames were collected continuously during the fractionation of the proteins (1 frame/sec). Frames collected before the void volume (0.6 – 1.5 ml) were averaged and subtracted from the signal of the elution profile to account for background scattering. Data reduction, subtraction, and averaging were performed using the software FOXTROT (SOLEIL). The scattering curves were analyzed using the package ATSAS,^44^ and reported as function of the angular momentum transfer q = 4π/λ sinθ, where 2θ is the scattering angle and λ the wavelength of the incident beam. Values of the cross-sectional radius of gyration were obtained with SCATTER.^45^

To model the dynamic behavior of SspH1 and explore its conformational space, we employed an Xplor-NIH protocol.^46^ Initially, the E2-UbBD (residues 623-697) and the LRR (residues 161-393) domains were randomly oriented relative to a fixed E3 domain (residues 405-622) by randomizing linker torsion angles (note: the LRR-E3 linker has a high B-factor). Subsequently, a high-temperature torsion-angle dynamic at 3000 K was followed by simulated annealing from 3000 K to 25 K in 12.5 K increments. The final step involved gradient minimization in torsion-angle space to generate the structure ensembles. Our calculations incorporated the experimental SAXS-derived force field, knowledge-based energy terms, such as torsion angle potential from conformational databases, and standard Xplor-NIH covalent and nonbonded energy terms. Each ensemble member was assigned a weight corresponding to 1/n, where n is the number of structures in the ensemble. Fifty equidistant points were used to simulate the scattering curve in a q interval of 0-0.3.

To avoid overfitting and determine the minimum number of conformers needed to accurately replicate the experimental SAXS data, we initially computed 100 ensembles with 1-6 conformers each (Figure S8B). The average χ2 rapidly decreased, plateauing for ensembles with three or more conformations. Subsequently, we calculated 2000 3-conformer ensembles and selected those with χ2 ≤ 1.5 for analysis (significance level α = 0.05, ensuring statistical significance in the association between the experimental data and our models). Distances between domain centers of mass and intra-domain angles were calculated using custom Python scripts. Envelope diameters and radii of gyration for each conformer were evaluated with the program Crysol (same reference of ATSAS).

### Nuclear Magnetic Resonance

NMR experiments were carried out using a Bruker AVANCE spectrometer operating at a proton nominal frequency of 800 MHz. Data were acquired with Topspin (Bruker), processed with NMRPipe,^47^ and analyzed by CCPNMR.^48^

For conjugated ubiquitin NMR, ^15^N isotope enriched UbMESNa was expressed and purified in M9 minimal medium using 1g/L of ^15^N-ammonium chloride as sole source of nitrogen. ^15^N UbMESNa was further reacted with glycine vinyl sulfone (EN300-1264982, Enamine) as described previously,^49^ to give ^15^N-ubiquitin vinyl sulfone. SspH1 constructs (10 µM) were reacted with 50 µM ^15^N UbVS for 2 hours before purification by gel filtration into 50 mM HEPES pH 7.5, 75 mM NaCl, 0.5 mM TCEP.

For ubiquitin titration NMR experiments, ^15^N isotope enriched ubiquitin was prepared by growing the bacteria in M9 minimal medium using 1g/L of ^15^N-ammonium chloride as sole source of nitrogen. ^15^N-ubiquitin was titrated with SspH1 constructs and measurements taken as described above. Chemical shifts changes for the backbone amide proton and nitrogen nuclei (ΔδNH) were calculated according to a procedure implemented in CCPNMR analysis.^48^

