## Supplementary Information for "Conformational dynamics of the bacterial E3 ligase SspH1"

**Contents**

Supplementary Figures 1-8
Supplementary Table1: X-ray data table
Supplementary Table 2: SAXS data table

A

```
IpaH1.4 ---------------------------------------------------------------------
IpaH9.8 ---------------------------------------------------------------------
SspH1   -MFNIRN--TQPSVSMQAIAGAAAPEASPEEIVWEKIQVFFPQENYEEAQQCLAELCHPARGMLPDHIS  66
SspH2   MPFHIGSGCLPATISNRRIYRIAWSDTPPEMSSWEKMKEFFCSTHQTEALECIWTICHPPAGTTREDVI  69

IpaH1.4 ------------------------------------------------------------MIKSTNIQ   8
IpaH9.8 ---------------------------------------------------------------------
SspH1   SQFARLKALTFPAWEENIQCNRDGINQFCILDAGSKEILSITLDDAGNYTVNCQGYSEAHDFIMDTEPG 135
SspH2   NRFELLRTLAYAGWEESIHSGQHGENYFCILDEDSQEILSVTLDDAGNYTVNCQGYSETHRLTLDTAQG 138

IpaH1.4 AIGSGIMHQINNVYSLTPLSLPMELTPSCNEFYLKTWSEWEKNGTPGEQRN--IAFNRLKICLQNQEAE  75
IpaH9.8 ------MLPINNNFSLPQNSFYNTI-SGTYADYFSAWDKWEKQALPGEERD--EAVSRLKECLINNSDE  60
SspH1   EECTEFAEGASG--T---SLRPATTVSQKAAEYDAVWSKWERDAPAGESPGRAAVVQEMRDCLNNGNPV 199
SspH2   EEGTGHAEGASG--TFRTSFLPATTAPQTPAEYDAVWSAWRRAAPAEESRGRAAVVQKMRACLNNGNAV 205

IpaH1.4 LNLSELDLKTLPDL-PPQITTLEIRKNLLTHLPDLPPMLKVIHAQFNQL-------------------- 123
IpaH9.8 LRLDRLNLSSLPDNLPAQITL--------------------LNVSYNQL--------------------  89
SspH1   LNVGASGLTTLPDRLPPHITTLVIPDNNLTSLPELPEGLRELEVSGNL-------------------QL 249
SspH2   LNVGESGLTTLPDCLPAHITTLVIPDNNLTSLPALPPELRTLEVSGNQLTSLPVLPPGLLELSIFSNPL 274

IpaH1.4 ESLPALPETLEELNAGDNKIKELPFLPENLTHLRVHNNRLHILPL--------------------LPPE 172
IpaH9.8 TNLPELPVTLKKLYSASNKLSELPVLPPALESLQVQHNELENLPA--------------------LPDS 138
SspH1   TSLPSLPQGLQKLWAYNNWLASLPTLPPGLGDLAVSNNQLTSLPE--------------------MPPA 298
SspH2   THLPALPSGLCKLWIFGNQLTSLPVLPPGLQELSVSDNQLASLPALPSELCKLWAYNNQLTSLPMLPSG 343

IpaH1.4 LKLLVVSGNRLDSIPPFPDKLEGLALANNFIEQLPEL------PFSMNR-------------------- 215
IpaH9.8 LLTMNISYNEIVSLPSLPQALKNLRATRNFLTELPAFSEGNN-PV-V-R-------------------- 184
SspH1   LRELRVSGNNLTSLPALPSGLQKLWAYNNRLTSLPEM------SPGLQELD------------------ 343
SspH2   LQELSVSDNQLASLPTLPSELYKLWAYNNRLTSLPAL------PSGLKELIVSGNRLTSLPVLPSELKE 406
                                               IpaH9.8 Amplifier loop

IpaH1.4 --------------------AVLMNNNLTTLPESVLRLAQNAFVNVAGNPLSGHTMRTLQQITTGPDYS 264
IpaH9.8 -------------------EYFFDRNQISHIPESILNLRNECSIHISDNPLSSHALPALQRLTSSPDYH 234
SspH1   ----------------------VSHNQLTRLPQSLTGLSSAARVYLDGNPLSVRTLQALRDII---GHS 387
SspH2   LMVSGNRLTSLPMLPSGLLSLSVYRNQLTRLPESLIHLSSETTVNLEGNPLSERTLQALREITSAPGYS 475

IpaH1.4 GPQIFFSMGNSATISAPEHSLADAVTAWFPENKQSD--VSQIWHAFEHEEHANTFSAFLDRLSDTVSAR 331
IpaH9.8 GPRIYFSMSDGQQN-TLHRPLADAVTAWFPENKQSD--VSQIWHAFEHEEHANTFSAFLDRLSDTVSAR 300
SspH1   GIRIHFDMAGPSVP-REARALHLAVADWLTSAREGEAAQADRWQAFGLEDNAAAFSLVLDRLRETENFK 455
SspH2   GPIIRFDMAGASAP-RETRALHLAAADWLVPAREGEPAPADRWHMFGQEDNADAFSLFLDRLSETENFI 543

IpaH1.4 NTSGFREQVAAWLEKLSASAELRQQSFAVAADATESCEDRVALTWNNLRKTLLVHQASEGLFDNDTGAL 400
IpaH9.8 NTSGFREQVAAWLEKLSASAELRQQSFAVAADATESCEDRVALTWNNLRKTLLVHQASEGLFDNDTGAL 369
SspH1   KDAGFKAQISSWLTQLAEDAALRAKTFAMATEATSTCEDRVTHALHQMNNVQLVHNAEKGEYDNNLQGL 524
SspH2   KDAGFKAQISSWLAQLAEDEALRANTFAMATEATSSCEDRVTFFLHQMKNVQLVHNAEKGQYDNDLAAL 612
                                  catalytic cysteine | catalytic acid

IpaH1.4 LSLGREMFRLEILEDIARDKVRTLHFVDEIEVYLAFQTMLAEKLQLSTAVKEMRFYGVSGVTANDLRTA 469
IpaH9.8 LSLGREMFRLEILEDIARDKVRTLHFVDEIEVYLAFQTMLAEKLQLSTAVKEMRFYGVSGVTANDLRTA 438
SspH1   VSTGREMFRLATLEQIAREKAGTLALVDDVEVYLAFQNKLKESLELTSVTSEMRFFDVSGVTVSDLQAA 593
SspH2   VATGREMFRLGKLEQIAREKVRTLALVDEIEVWLAYQNKLKKSLGLTSVTSEMRFFDVSGVTVTDLQDA 681
                                  catalytic base

IpaH1.4 EAMVRSREENEFTDWFSLWGPWHAVLKRTEADRWAQAEEQKYEMLENEYSQRVADRLKASGLSGDADAE 538
IpaH9.8 EAMVRSREENEFTDWFSLWGPWHAVLKRTEADRWAQAEEQKYEMLENEYPQRVADRLKASGLSGDADAE 507
SspH1   ELQVKTAENSGFSKWILQWGPLHSVLERKVPERFNALREKQISDYEDTYRKLYDEVLKSSGLVDDTDAE 662
SspH2   ELQVKAAEKSEFREWILQWGPLHRVLERKAPERVNALREKQISDYEETYRMLSDTELRPSGLVGNTDAE 750
                                                        E2~Ub binding

IpaH1.4 REAGAQVMRETEQQIYR---QLTDEVLALRLSENGSNHIA-                             575
IpaH9.8 REAGAQVMRETEQQIYR---QLTDEVLALRLPENGSQLHHS                             545
SspH1   RTIGVSAMDSAKKEFLDGLRALVDEVLGSYLTARWRLN---                             700
SspH2   RTIGARAMESAKKTFLDGLRPLVEEMLGSYLNVQWRRN---                             788
        E2~Ub binding
```

B

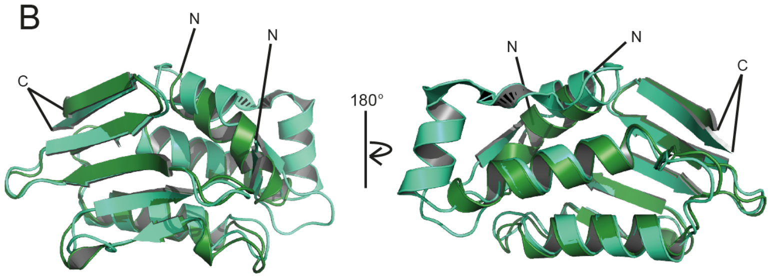


C

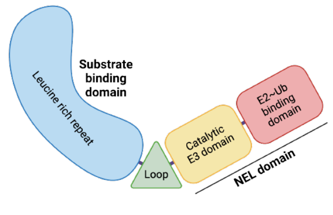


**Supplementary Figure 1** NEL protein sequence alignment.

A) Alignment of sequences of IpaH1.4, IpaH9.8, SspH1 and SspH2 using Clustal Omega, with SspH1 secondary structure shown above. N-terminal domains of SspH1 and SspH2 as predicted by AlphaFold2 are highlighted in green and turquoise respectively. SspH1 domains are highlighted in blue (LRR), green (flexible loop between LRR and NEL domains), yellow (E3 catalytic domain) and light red (E2-Ub binding domain). The catalytic cysteines, catalytic acids and bases are highlighted in red, and critical conserved residues highlighted in orange.[1] Grey residues represent those critical for E2-Ub binding.[1] B) Aligned AF2 predictions of N-terminal domains of SspH1 (green) and SspH2 (turquoise). C) Domain structure of NEL bacterial E3 ligases, with colours corresponding to domains highlighted in the sequence alignment.

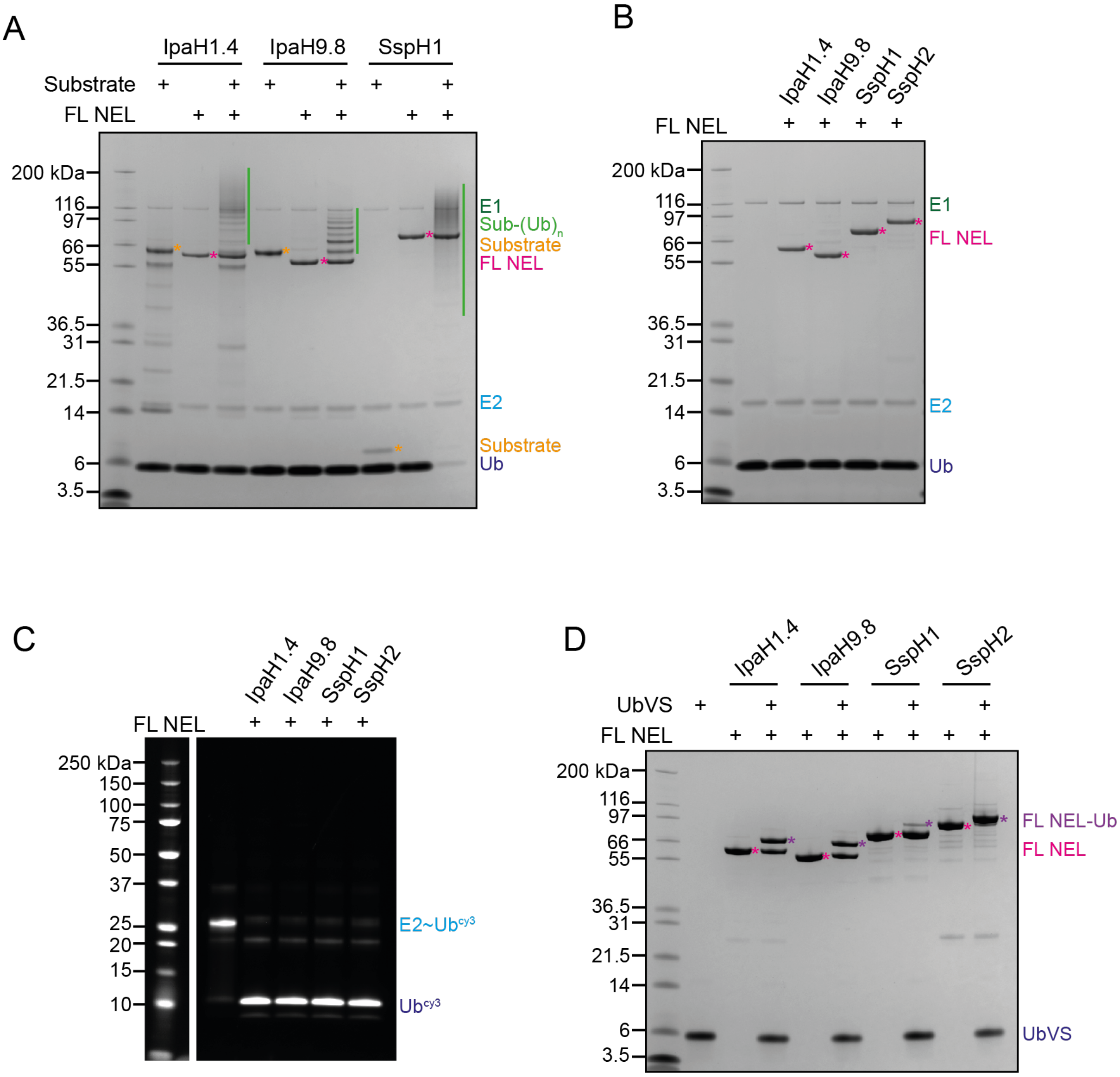


**Supplementary Figure 2** Full length (FL) construct assays.

A) Substrate ubiquitination assay with FL constructs of IpaH1.4 (substrate LUBAC), IpaH9.8 (substrate GBP1) and SspH1 (substrate HR1b PKN1). Performed with 0.1 µM UBA1 (E1), 2 µM UbcH5A (E2), 1.0 µM FL NEL (E3), 20 µM ubiquitin, 10 mM ATP at RT for 30 minutes with either 1 µM (LUBAC, GBP1) or 2 µM (HR1b PKN1) substrate.

B) Auto-ubiquitination assay with FL constructs of IpaH1.4, IpaH9.8, SspH1 and SspH2. Performed with 0.1 µM UBA1 (E1), 2 µM UbcH5A (E2), 1.0 µM FL NEL (E3), 20 µM ubiquitin, 10 mM ATP at RT for 30 minutes.

C) E2~Ub discharge assay with FL constructs of IpaH1.4, IpaH9.8, SspH1 and SspH2. Performed with 1 µM UbcH5A~Ub-cy3 (E2~$Ub^{cy3}$), 0.1 µM FL NEL (E3) at RT for 20 minutes.

D) Ubiquitin-loading assay with FL constructs of IpaH1.4, IpaH9.8, SspH1 and SspH2. Performed with 20 µM UbVS, 5 µM FL NEL (E3) at RT for 2 hours.

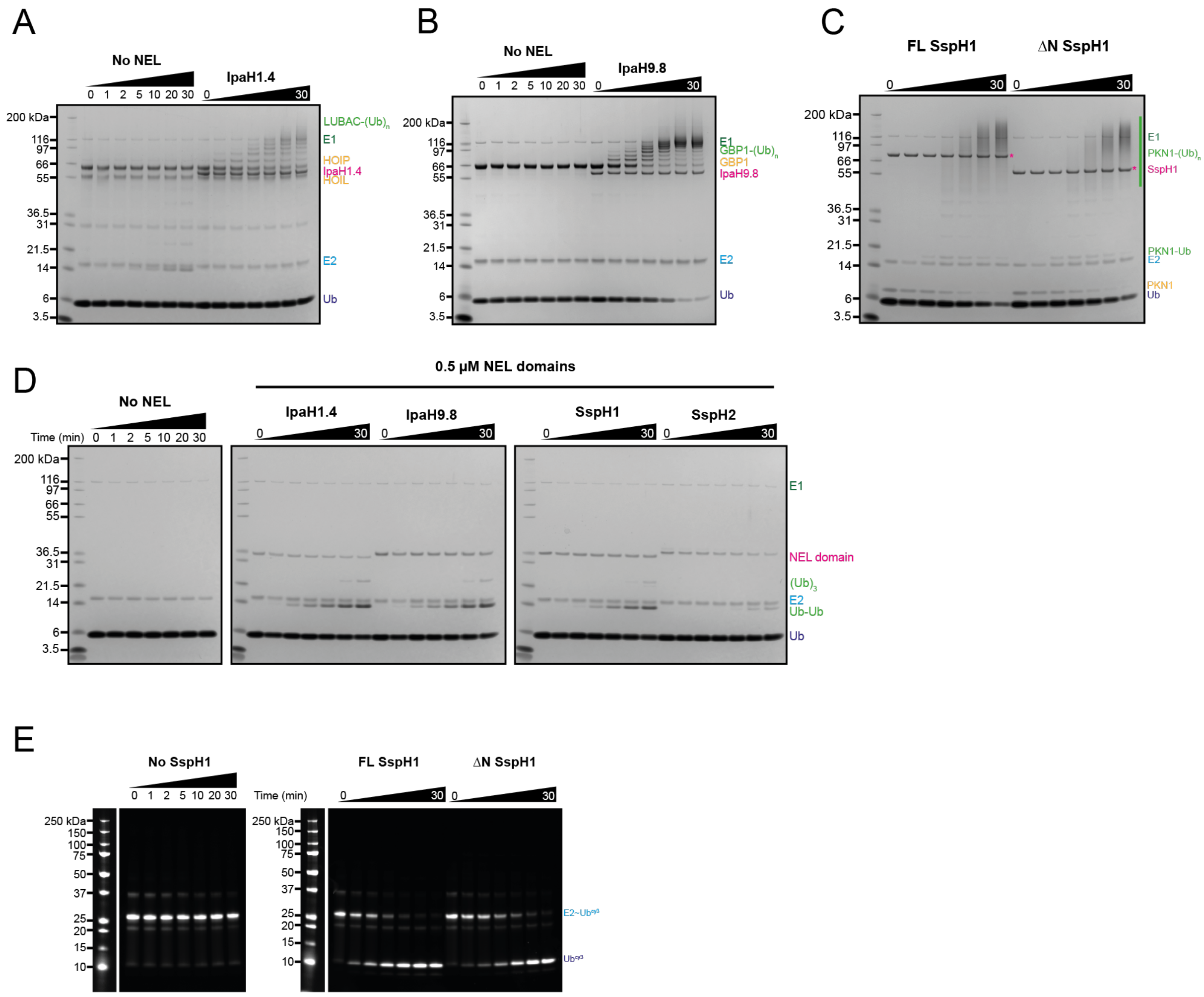


**Supplementary Figure 3** Time course assays.

A) Substrate ubiquitination assay with ΔN IpaH1.4 with LUBAC substrate. Performed with 0.1 µM UBA1 (E1), 2 µM UbcH5A (E2), 1 µM IpaH1.4 (E3), 1 µM LUBAC, 20 µM ubiquitin, 10 mM ATP at RT for 0-30 minutes.

B) Substrate ubiquitination assay with ΔN IpaH9.8 with GBP1 substrate. Performed with 0.1 µM UBA1 (E1), 2 µM UbcH5A (E2), 1.0 µM IpaH9.8 (E3), 1 µM GBP1, 20 µM ubiquitin, 10 mM ATP at RT for 0-30 minutes.

C) Substrate ubiquitination assay with FL SspH1 and ΔN SspH1 with HR1b PKN1 substrate. Performed with 0.1 µM UBA1 (E1), 2 µM UbcH5A (E2), 1.0 µM SspH1 (E3), 2 µM HR1b PKN1, 20 µM ubiquitin, 10 mM ATP at RT for 0-60 minutes.

D) Auto-ubiquitination assay with NEL domains of IpaH1.4, IpaH9.8, SspH1 and SspH2. Performed with 0.1 µM UBA1 (E1), 2 µM UbcH5A (E2), 0.5 µM NEL domain (E3), 20 µM ubiquitin, 10 mM ATP at RT for 0-30 minutes.

E) E2~Ub discharge assay with FL SspH1 and ΔN SspH1. Performed with 1 µM UbcH5A~Ub-cy3 (E2~Ub$^{cy3}$), 0.1 µM SspH1 (E3) at RT for 0-30 minutes.

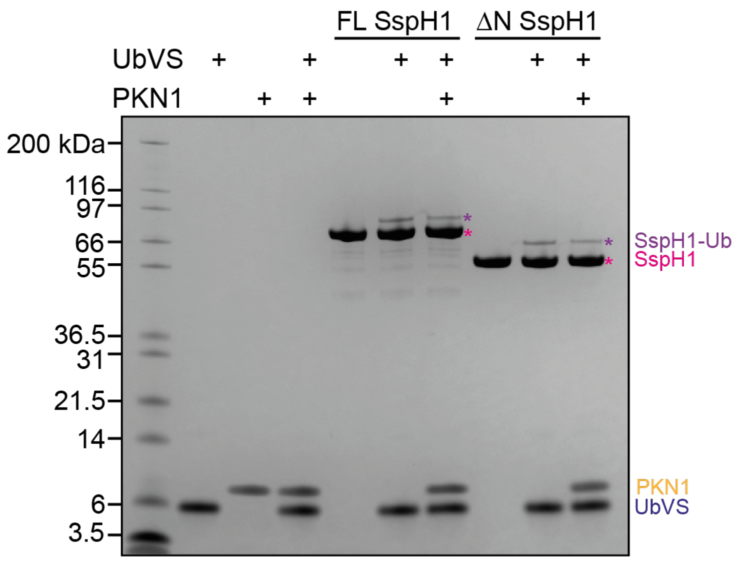


**Supplementary Figure 4** Loading of SspH1 ΔN and FL constructs with UbVS with and without PKN1.

A) Ubiquitin-loading assay with FL and ΔN SspH1, with and without HR1b PKN1. Performed with 20 μM UbVS, 5 μM FL NEL (E3), 10 μM HR1b PKN1 at RT for 2 hours.

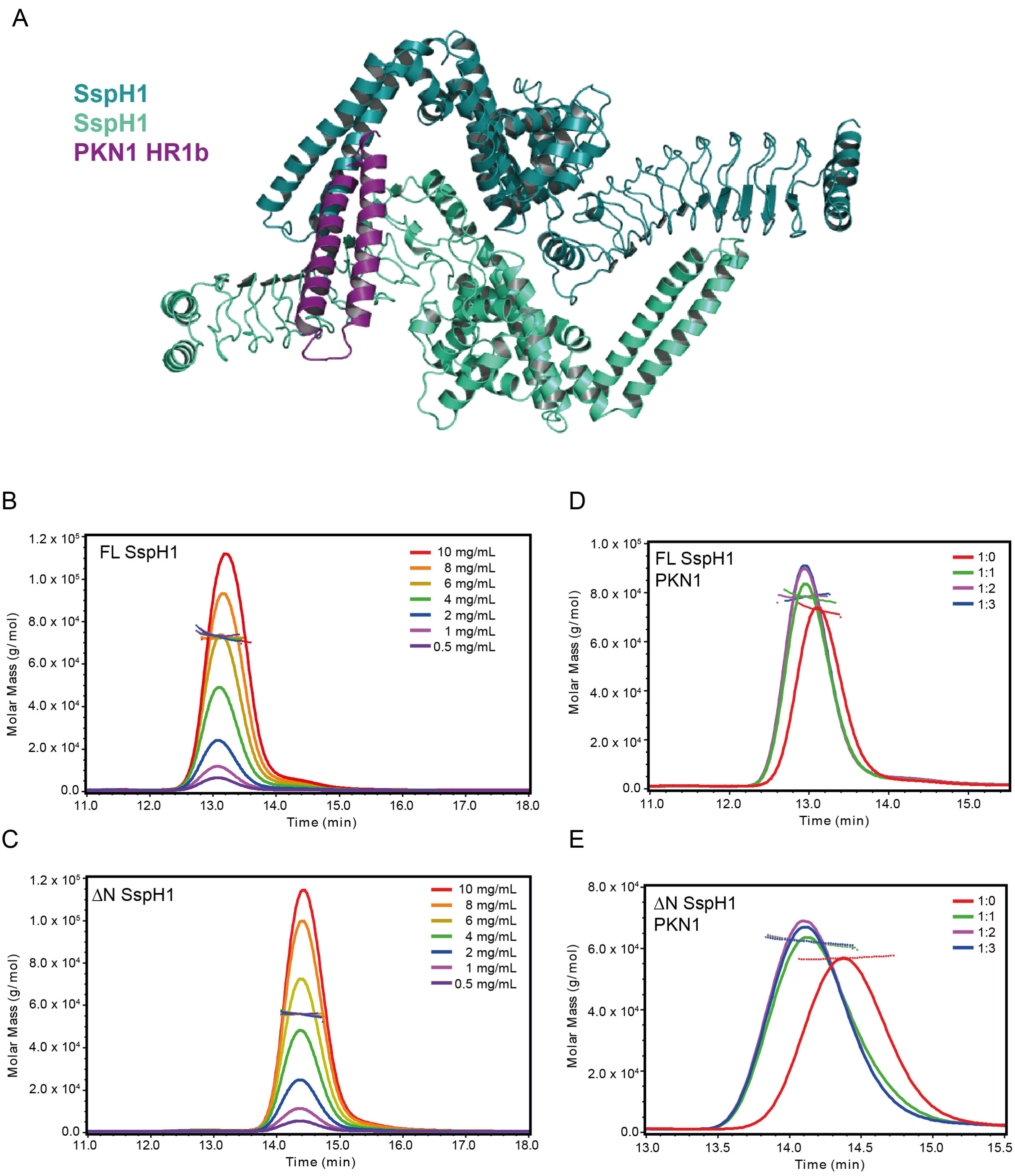


**Supplementary Figure 5** SspH1 is a monomer.

A) Symmetry related molecules of ΔNSspH1 (teal and turquoise) aligned to PKN1 HR1b (purple) in complex with SspH1 LRR domain (PDB 4NKG); SEC-MALLS of B) FL SspH1; C) ΔNSspH1; D) FL SSpH1 titrated with PKN1 HR1b; E) ΔNSspH1 titrated with PKN1 HR1b.

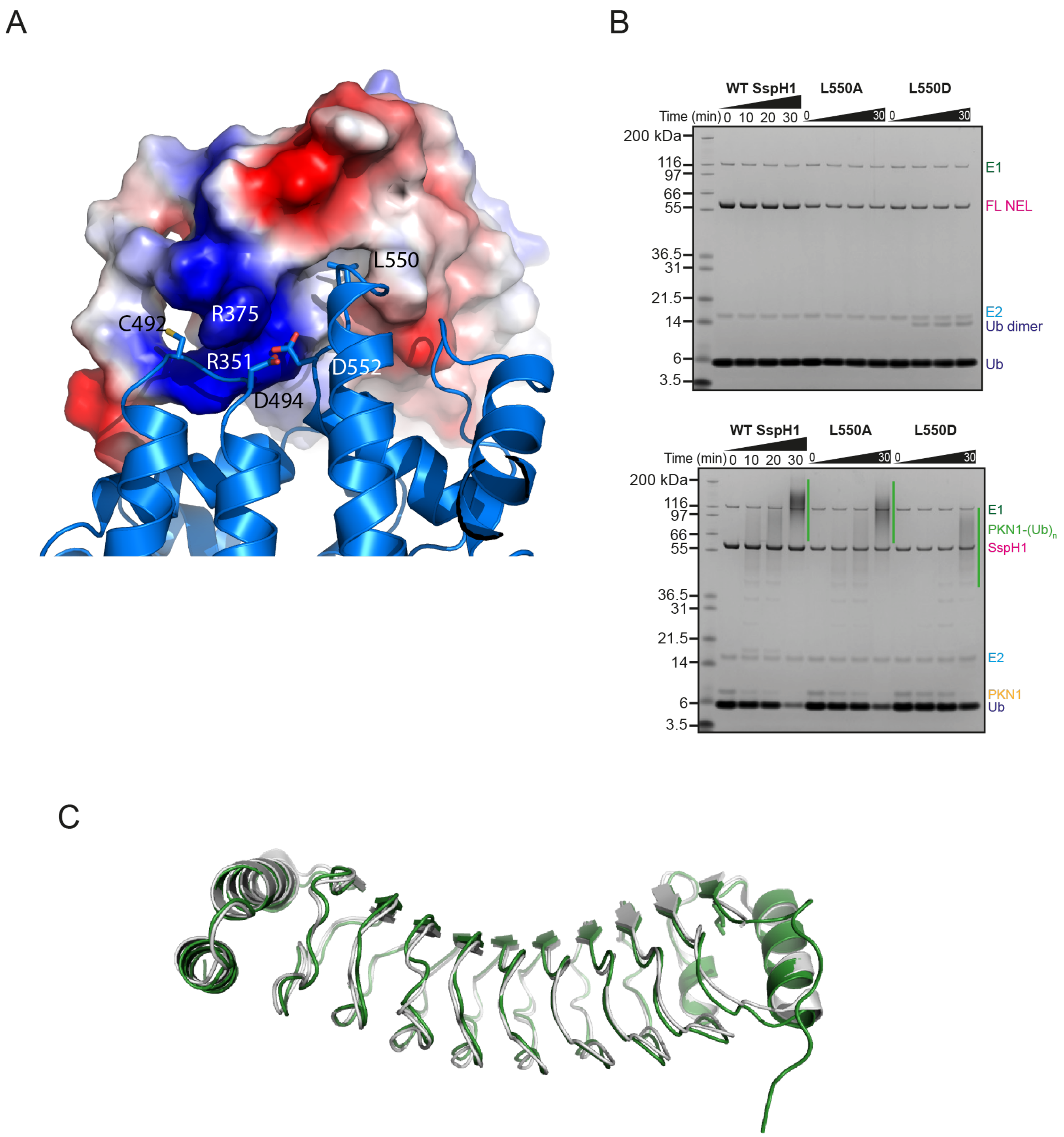


**Supplementary Figure 6** SspH1 mutant assays.

A) Close up view of the interface between LRR and NEL domain, with the NEL in surface representation coloured according to electrostatic potential and the NEL in cartoon representation. L550 points into a hydrophobic pocket formed by the LRR, whereas R351 and R375 from the LRR contact acidic residues of the NEL domain.

B) Substrate ubiquitination assay with ΔNSspH1 mutants L550A and L550D. Performed with 0.1 μM UBA1 (E1), 2 μM UbcH5A (E2), 1.0 μM SspH1 (E3), 2 μM HR1b PKN1, 20 μM ubiquitin, 10 mM ATP at RT for 0-30 minutes.

C) Overlap of the LRR domains of the SspH1 structure presented here in green and the LRR domain structure bound to PKN1 HR1b (PDB: 4NKG) in grey[2].

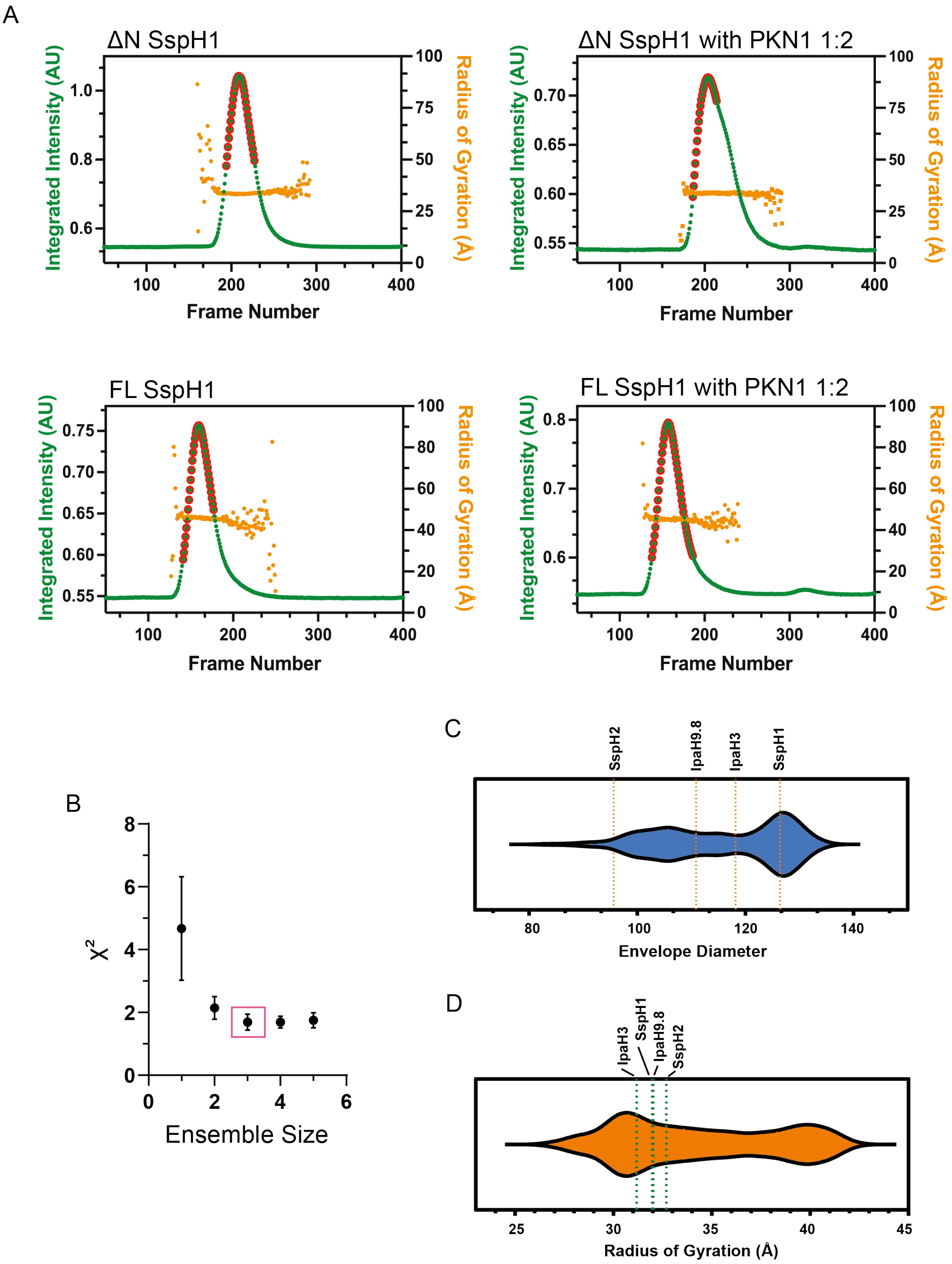


**Supplementary Figure 7** SEC-SAXS analysis and Xplor-NIH modelling.
A) Integrated intensities of recorded SAXS as a function of frames recorded off the size-exclusion column. The orange points are the values of the radius of gyration of the eluting species along the gel-filtration peak. Highlighted in red are the averaged frames for the final scatting curve. B) SAXS-based Xplor-NIH calculated ensemble $\chi^2$ values versus the size of the ensemble. The lowest $\chi^2$ values are achieved with a minimum ensemble size made of 3 conformers. C) Envelope diameter and radius of gyration distributions of the calculated ensemble conformers with $\chi^2$ values smaller than 1.5. The values for the available full-length structures of SspH2 (PDB 3G06), IpaH9.8 (PDB 6LOL), IpaH3 (PDB 3CVR) are reported with the values obtained for our crystal structure.

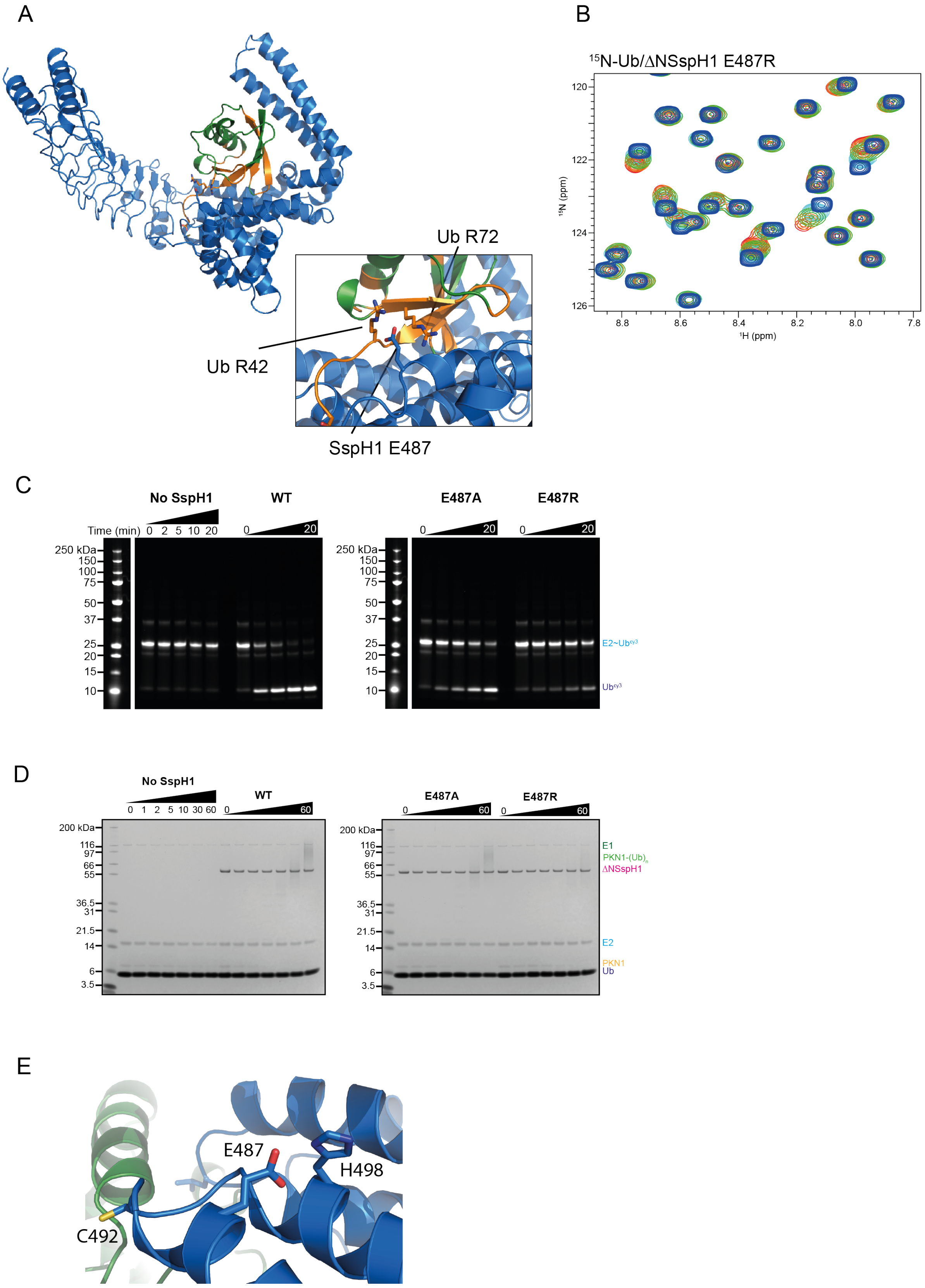


**Supplementary Figure 8** Analysis of the SspH1-ubiquitin interaction.

A) AF3 prediction of ubiquitin (green) bound to ΔNSspH1 (blue), with perturbed ubiquitin residues shown in orange. Insert: SspH1 E487 is predicted to form interactions with perturbed residues R72 and R42 of ubiquitin.
B) Details of the titration of $^{15}$N-labeled ubiquitin titrated with ΔNSspH1 E487R. Spectra at different ligand concentrations are plotted at the same contour level.
C) E2~Ub discharge assay with ΔNSspH1 E487A and E487R mutants. Performed with 1 μM UbcH5A~Ub-cy3 (E2~Ub$^{cy3}$), 0.1 μM SspH1 (E3) at RT for 0-30 minutes.
D) Substrate ubiquitination assay with ΔNSspH1 E487A and E487R mutants with HR1b PKN1 substrate. Performed with 0.1 μM UBA1 (E1), 2 μM UbcH5A (E2), 1.0 μM SspH1 (E3), 2 μM HR1b PKN1, 20 μM ubiquitin, 10 mM ATP at RT for 0-30 minutes.
E) Close up of the interaction between E487 and H498 which are positioned in the helices either side of the loop containing the catalytic cysteine.

**Supplementary Table 1. Data collection and refinement statistics.**

| | **SspH1 161-700** |
|---|---|
| **Wavelength (Å)** | 0.62 |
| **Resolution range** | 58.31 - 2.9 (3.0 - 2.9) |
| **Space group** | P 6 2 2 |
| **Unit cell** | 170.8 170.8 94.8 90 90 120 |
| **Total reflections** | 735501 (73343) |
| **Unique reflections** | 18583 (1811) |
| **Multiplicity** | 39.6 (40.5) |
| **Completeness (%)** | 99.8 (99.6) |
| **Mean I/sigma(I)** | 6.8 (0.4) |
| **Wilson B-factor** | 74.61 |
| **R-merge** | 0.4992 (6.506) |
| **R-meas** | 0.5056 (6.588) |
| **R-pim** | 0.07988 (1.03) |
| **CC1/2** | 0.996 (0.416) |
| **CC*** | 0.999 (0.767) |
| **Reflections used in refinement** | 18544 (1804) |
| **Reflections used for R-free** | 900 (105) |
| **R-work** | 0.2595 (0.3720) |
| **R-free** | 0.3158 (0.4481) |
| **CC(work)** | 0.943 (0.630) |
| **CC(free)** | 0.832 (0.497) |
| **Number of non-hydrogen atoms** | 4247 |
| **macromolecules** | 4193 |
| **ligands** | 14 |
| **solvent** | 44 |
| **Protein residues** | 540 |
| **RMS(bonds)** | 0.001 |
| **RMS(angles)** | 0.36 |
| **Ramachandran favored (%)** | 98.88 |
| **Ramachandran allowed (%)** | 1.12 |
| **Ramachandran outliers (%)** | 0.00 |
| **Rotamer outliers (%)** | 0.66 |
| **Clashscore** | 18.89 |
| **Average B-factor** | 78.93 |
| **macromolecules** | 79.16 |
| **ligands** | 59.55 |
| **solvent** | 61.58 |

Statistics for the highest-resolution shell are shown in parentheses.

### Supplementary Table 2: SAXS parameters and data analysis software.

**Supplementary Table 1: SAXS parameters.**

| **Data collection** | | | | |
|---|---|---|---|---|
| Beamline | SWING at Soleil | | | |
| q range ($Å^{-1}$) | 0.0082–0.66 | | | |
| Detector | EigerX4M in vacuum | | | |
| Column | Bio-SEC 3 Agilent 300 Å | | | |
| Flow rate (ml/min) | 0.3 | | | |
| Temperature (°C) | 15 | | | |
| **Samples details** | **ΔNSspH1** | **ΔNSspH1/PKN1** | **SspH1** | **SspH1/PKN1** |
| Sample volume (µl) | 60 | 60 | 60 | 60 |
| Sample concentration (mg/ml) | 10.0 | 11.5 | 8.0 | 9.0 |
| **Structural parameters** | | | | |
| **Reciprocal Space** | | | | |
| Rg (Å) Guinier | 34.5 | 35.1 | 48.2 | 47.7 |
| I(0) ($cm^{-1}$) | 0.11 | 0.09057 | 0.01101 | 0.02078 |
| qRg limit | 1.05 | 1.04 | 1.16 | 0.97 |
| **Real Space** | | | | |
| Rg (Å) P(R) | 34.6 ± 0.01 | 35.2 ± 0.05 | 48.5 ± 0.1 | 47.8 ± 0.08 |
| I(0) ($cm^{-1}$) | 0.11000 ± 0.00003 | 0.09058 ± 0.00009 | 0.01102 ± 0.00001 | 0.02078 ± 0.00001 |
| Rc (Å) | 19.6 | 20.7 | 18.6 | 20.2 |
| Dmax (Å) | 114 | 124 | 189 | 198 |
| Porod volume ($Å^3$) | 86317 | 104099 | 113536 | 143322 |
| **Molecular mass determination** | | | | |
| Theoretical MW (kDa) | 59.5 | 68.8 | 77.1 | 86.4 |
| DATPOROD MW (kDa) (Vp/1.6) | 53.9 | 65.1 | 71.0 | 89.6 |
| SAXS MoW2 (q = 0.3 $Å^{-1}$) | 61.9 | 70.1 | 82.9 | 91.9 |

| **Data analysis software** | |
|---|---|
| Primary Data Reduction | Foxtrot |
| Data processing | Primus & Scatter |
| Computation of scattering intensities | Crysol |
| Structural modelling | Xplor-NIH |
| 3D graphics representation | Pymol |
